# Trait-based responses to land use and canopy dynamics modify long-term diversity changes in forest understories

**DOI:** 10.1101/2020.04.30.069708

**Authors:** Konsta Happonen, Lauralotta Muurinen, Risto Virtanen, Eero Kaakinen, John-Arvid Grytnes, Elina Kaarlejärvi, Philippe Parisot, Matias Wolff, Tuija Maliniemi

## Abstract

**Aim:** Land use is the foremost cause of global biodiversity decline, but species do not respond equally to land-use practices. Instead, it is suggested that responses vary with species traits, but long-term data on the trait-mediated effects of land-use on communities is scarce. Here we study how forest understory communities have been affected by two land-use practices during 4–5 decades, and whether changes in plant diversity are related to changes in functional composition.

**Location:** Finland

**Time period:** 1968–2019

**Major taxa studied:** Vascular plants

**Methods:** We resurveyed 245 vegetation plots in boreal herb-rich forest understories, and used hierarchical Bayesian linear models to relate changes in diversity, species composition, average plant size, and leaf economic traits to reindeer abundance, forest management intensity, and changes in climate, canopy cover and composition.

**Results:** Intensively managed forests decreased in species richness and had increased turnover, but management did not affect functional composition. Increased reindeer densities corresponded with increased leaf dry matter content, evenness and diversity, and decreased height and specific leaf area. Successional development in the canopy was associated with increased specific leaf area and decreased leaf dry matter content and height in the understory over the study period. Effects of reindeer abundance and canopy density on diversity were partially mediated by vegetation height, which had a negative relationship with evenness. Observed changes in climate had no discernible effect on any variable.

**Main conclusions:** Functional traits are useful in connecting vegetation changes to the mechanisms that drive them, and provide unique information compared to turnover and diversity metrics. These trait-dependent selection effects could inform whether species benefit or suffer from land use changes and explain observed biodiversity changes under global change.

## Introduction

There is an urgent need to develop better tools for monitoring biodiversity change, as the most commonly used metrics, species richness and compositional turnover, do not capture its full extent (Blowes et al., 2019; Hillebrand et al., 2018). One way to move towards understanding the processes behind biodiversity change is to establish links between the functional composition of communities and potential selective forces, such as land-use change. Changes in average functional traits can be directly informative of changing selection pressures on communities and resulting ecosystem consequences (Lavorel & Garnier, 2002; Violle et al., 2007). That is, if we knew how the functional composition of communities responded to anthropogenic pressures, we could use traits to monitor the effects of human actions on natural systems, and perhaps predict the resulting changes in ecosystem properties.

The above-ground traits of vascular plant species and communities have been found to vary on at least two important and independent axes: plant size and the leaf economics spectrum (LES). Size-related traits determine competitive hierarchies in relation to light, while leaf economic traits describe, for example, leaf construction costs and maximum photosynthetic output (Bruelheide et al., 2018; Díaz et al., 2016). Plant size is often described with vegetative height, and the LES with traits such as specific leaf area (SLA) and leaf dry matter content (LDMC) (van der Plas et al., 2020). When external drivers such as land-use change affect the optimal allocation of resources to vertical growth and to leaves, it is also expected to exert directional selection pressure on the composition of plant communities, which should manifest as changes in average functional traits.

Climate has traditionally been seen as the dominant driver of boreal biodiversity, while the impacts of other drivers such as large herbivores have been underestimated; yet they play a prominent role in community assembly (Pausas & Bond, 2018). Large herbivores have strong impacts on vegetation that include favoring traits that decrease susceptibility to grazing, such as small size and unpalatable leaves (e.g. high LDMC and low SLA, Díaz et al., 2007). Nowadays wild megafauna has been partly replaced by human livestock (Hempson, Archibald, & Bond, 2017), and humans regulate herbivore densities. In many parts of Eurasia and Northern America reindeer/caribou (*Rangifer tarandus*) is herded as semi-domesticated free-ranging livestock. Reindeer grazing is known to cause declines in total plant biomass and in favoured forage species such as large herbs and deciduous shrubs in boreal and tundra environments (Bråthen & Oksanen, 2001; Olofsson, Moen, & Östlund, 2010; Sundqvist et al., 2019). Evidence of the impact of large herbivores on plant traits exists from other systems (Cingolani, Cabido, Gurvich, Renison, & Díaz, 2007; Díaz et al., 2007), but is very limited from the boreal zone and especially from its forested systems.

One third of global forested area belongs to the boreal zone, and a majority of it is used for industrial wood production (Gauthier, Bernier, Kuuluvainen, Shvidenko, & Schepaschenko, 2015). Management effects on ecosystems can lag decades behind changes in management practices (Muurinen, Oksanen, Vanha-Majamaa, & Virtanen, 2019), underlining the importance of long-term resurvey data for detecting biodiversity changes. Even though boreal forests are estimated to be highly sensitive to modern climate change (IPCC, 2018), the species compositions of both canopies and understories have shown relatively small long-term responses to climate change compared to land-use (Danneyrolles et al., 2019; Tonteri et al., 2016). A possible reason for the weak climate change responses is that closed forest canopies can buffer microclimatic conditions in the understory against macroclimatic warming (De Frenne et al., 2019). Trends in canopy cover can be affected by land-use histories, as disturbed tree-layers densen naturally when left alone, while new disturbances, such as logging, can increase temperature and light availability in the understory (Tonteri et al., 2016). Changes in light availability should be reflected in SLA, which increases in response to shade (Dahlgren, Eriksson, Bolmgren, Strindell, & Ehrlén, 2006). A recent resurvey from European temperate forests shows increasing canopy shade to be the primary cause of changes in functional composition of herb-layers in these systems (Depauw et al., 2020), but the generality of this finding in other biomes remains untested.

The monitoring of average functional traits alongside diversity is also important because diversity is not independent of functional composition. It has long been known that high productivity can lead to decreased diversity (Grime, 1973). The unimodal productivity-diversity relationship is in part a mechanistic consequence of productivity increasing average species size, since communities composed of larger species have room for fewer plant individuals and therefore exhibit decreased diversity (Oksanen, 1996). Similar hypotheses regarding size-diversity relationships have been put forward and tested for other taxa as well (Siemann, Tilman, & Haarstad, 1996). Thus, in systems where human actions modify the selective landscape in relation to species size, anthropogenic disturbances are expected to also affect local diversity. For example, as discussed above, increased grazing pressure by livestock should proportionately favor smaller species and thus increase diversity (Jia et al., 2018).

Because the processes that drive changes in plant communities may be spatially structured, changes in community properties may be as well. For example, diversity changes might be positive in some areas and negative in others, resulting in zero average change (Vellend et al., 2013). This needs to be taken into account when interpreting average changes, lest we fall victim to the ecological fallacy of assuming that all members of a population behave in the same way as the population on average (Robinson, 1950).

Here we study the temporal and spatial variation in diversity and community composition by analysing the long-term effects of two main land-use types (reindeer husbandry and forestry) and changes in canopies (canopy SLA and cover) on boreal herb-rich forest understories. By comparing diversity-, dissimilarity-, and functional trait -based approaches using hierarchical Bayesian regression modelling, we seek answers to four questions: 1) have species diversity, functional composition (measured as average height, SLA and LDMC), and species composition changed in boreal herb-rich forests over 40–50 years, 2) are there spatial differences in these changes 3) are the changes related to changes in semi-domesticated reindeer abundance, forest management intensity, or changes in canopy SLA and cover, and finally 4) are the changes in functional composition correlated with changes in diversity across both time and space, hinting at processes that link traits to diversity?

## Materials and methods

### Vegetation data and resampling

We conducted vegetation resurvey in boreal herb-rich forests in Northern Finland (Fig. 1). The resurveyed herb-rich forest sites were originally surveyed in 1968-75 by Eero Kaakinen (Kaakinen, 1971, 1974; Unpublished). The purpose of the original surveys was to identify phytosociological species associations in mature herb-rich forests. For this reason, the surveys were conducted in relatively undisturbed habitats. These forests harbour higher species richness than the surrounding less fertile and homogenous boreal forests (Maliniemi, Happonen, & Virtanen, 2019), are an important habitat for many threatened species (Kouki et al., 2018), and are thus important to monitor. They occur typically as small patches and cover only a fraction of the forested area in northern Scandinavia reflecting the scattered distribution of calcareous bedrock and soil. The field layer is dominated by herbs such as *Geranium sylvaticum* and *Filipendula ulmaria*, ferns such as *Gymnocarpium dryopteris* and *Athyrium filix-femina*, and graminoids such as *Milium effusum* and *Elymus caninus*. Occasionally, they also host species typical for less fertile boreal forests, such as the dwarf-shrubs *Vaccinium myrtillus* and *Vaccinium vitis-idaea*. The three most abundant tree species *Picea abies*, *Alnus incana*, and *Betula pubescens* formed more than 80% of tree cover in our study plots, during both the original survey and the resurvey.

**Fig. 1:**
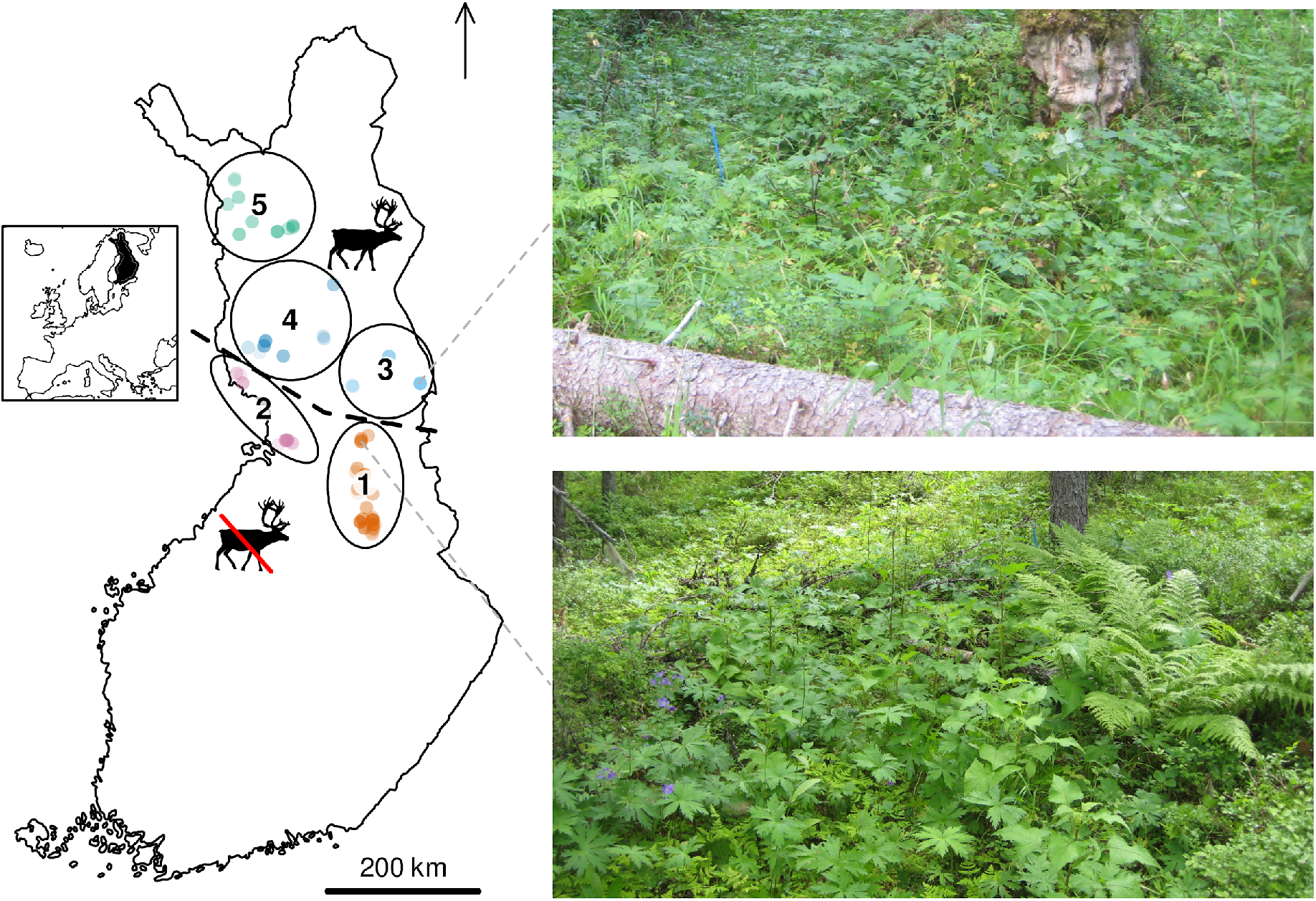
Map of the study area. The circles depict the biogeographical district (BGD) division used in this study. The line in the middle is the southern edge of the reindeer herding area. The images on the right depict typical vegetation in reindeer-grazed (upper) and ungrazed (lower) herb-rich forests. Basemap CC-BY National Land Survey of Finland.

Using detailed notes on the locations of the original sites, we relocated and resurveyed a majority of the original 336 sites in 2013-2019. Most surveys were done in teams of ≥2 investigators. We omitted sites which lacked detailed enough descriptions of location, and some sites that were located very far from other sites. The final number of plots, after applying exclusion criteria described below, was 245. Relocation error is inherent in vegetation resurveys (Kapfer et al., 2017), but was minimized as much as possible by the relocation help of the original surveyor, the patchy distribution of the studied vegetation type, and the fact that herb-rich forests are easily told apart from the boreal forest matrix.

This study thus applies all of the measures listed by Verheyen et al. (2018) for minimizing observer and relocation error. During both the original survey and the resurvey, species composition was estimated from 5 m x 5 m quadrats for the field layer and from 10 m x 10 m quadrats for shrub and canopy layers. The smaller quadrats were nested in the larger quadrats. The abundances of field and shrub-layer species were estimated as absolute percentage covers, and the covers of tree-layer species were estimated as relative covers, i.e. they always summed to 100%. In addition, total canopy cover was estimated visually on a three-point ordinal scale (0–30%, 31–70%, 71–100%). In this study, we focus only on the vascular plants of the field and tree layers. Species in the shrub layer (woody species generally over 0.5 m but under 2 m tall) were excluded from analyses because they were measured at a different plot size compared to the field layer, and was deemed justified because the cover of such species was low (~2% on average). For species list and trait values, see Table S1.1 in Appendix S1 in Supporting Information.

### Environmental changes

We estimated forest management intensity during the resurvey based on the protocol used for the assessment of threatened habitat types in Finland (Kouki et al., 2018, p. 180). This protocol assigns sites on an ordinal scale of habitat quality ranging from zero to four, describing absence of forest management impacts. Here, we inverted the scale to describe management intensity. We also omitted one site that was destroyed by deforestation and land conversion to arrive at a management impact variable ranging from one to four. The management intensity assessment criteria include the structure of the canopy and vegetation layers, disturbance of the soil, and the amount of deadwood, and are described in Table S2.2 (See Appendix S2). These criteria are applied in a one-out-all-out manner, meaning that if any of the criteria for a more intensive management class were fulfilled, the site was assigned in that class even if it would have been a less intensively managed site based on all other criteria. The criteria are independent of the other variables used in this study, meaning that species composition and diversity were not used as measures of management intensity. Signs of forest management were present in *ca.* 70% of the resurveyed plots, whereas the rest *ca.* 30% were deemed unmanaged (Fig. S2.1). Our observations in the field (including old stumps, plantations of c. 30-year-old trees, and old ditches) confirm that forest management was not concentrated near the second resampling, but was spread over a longer time period.

Reindeer are the major large grazer in our study area. In Finland, semi-domesticated reindeer occur only inside a designated reindeer herding area. Data on reindeer abundances in the different reindeer herding districts were provided by Natural Resource Institute Finland. Reindeer densities in the herding area had increased *ca.* 40% during the study period, from 1 to 1.4 reindeer per km^−2^ (Fig. S2.2). We calculated 10-year average reindeer densities prior to each sampling time to be used as a predictor of vegetation changes. The population sizes of other large herbivores in the study area, as inferred from hunting statistics, are negligible (Natural Resources Institute Finland, 2020, Fig. S2.1), with the exception of the moose (*Alces alces*). Although the moose population in Finland has grown due to changes in forestry practices and hunting, the growth has been rather uniform across the country (Nygrén, 2009). Moreover, herbaceous field layer vegetation constitutes less than 5% of moose summer diets (Hjeljord, Hovik, & Pedersen, 1990). Thus, while moose density changes might have affected forest regeneration and shrub layer composition, the direct effects of increased moose densities should be rather small and uniform across the study area. Contrarily, fertile herb-rich forests are a key foraging habitat for reindeer during summer, and we expect that changes in reindeer abundances over time can strongly affect trait composition and diversity of plant communities.

We included climate change in our analyses to test its potential effect on vegetation. Ten-year average minimum and maximum summer temperatures in our study plots have increased by 0.6℃ and 0.8℃ in 45 years (1970–2015, Fig. S2.4), and changes experienced by the field layer will have been further buffered by the canopy layer (De Frenne et al., 2019). To avoid choosing between highly correlated maximum and minimum temperatures and implying that these interpolated variables carry more information than they do, we chose to explain vegetation responses to climate using 10-year average vapor pressure deficit (VPD) during the summer months from the Terra Climate dataset (Abatzoglou, Dobrowski, Parks, & Hegewisch, 2018). VPD calculated this way has a correlation of 0.9 and 1.0 with average minimum and maximum summer temperatures from the same dataset, and has the added benefit of being defined in terms of average climatic conditions instead of thermal extremes. Averages were calculated from years 1961–70 for the original sample and years 2006–15 for the resample.

### Community properties

All data processing and analyses were done in R (version 4.0.5, R Core Team, 2019).

As species diversity measures, we calculated species richness, Shannon diversity, and species evenness. By Shannon diversity we mean the so-called true diversity of order 1, which is a unit conversion of the Shannon diversity index from entropy to species (Jost, 2006). As species evenness, we used Pielou’s J, or the ratio of Shannon entropy to log-transformed species richness, which is a measure of relative evenness ranging from zero to one (Jost, 2010).

We calculated community weighted means (CWM) of log-transformed trait values for one size-structural trait and two leaf economic traits: vegetative height, specific leaf area (SLA), and leaf dry-matter content (LDMC). Trait values were log-transformed because generally, traits follow a log-normal distribution (Bjorkman et al., 2018). Species absolute covers were transformed to relative covers and used as weights. Trait values were retrieved from the TRY database (Kattge et al., 2020, version 5, 2011) and the LEDA database (Kleyer et al., 2008), and supplemented with our own measurements for common species that were not found from the databases. In four plots, some of the traits were available for less than 80% of total cover. These plots were excluded from analyses. In the remaining plots, trait values were available for >99.5% of total cover for all traits. References to the original trait datasets are listed in Appendix S3.

We also calculated a CWM for the SLA of the canopy layer. SLA was not available for the non-native *Larix sibirica* (present on one site), for which we used the SLA of *Larix decidua* instead.

Summary statistics of the plant community properties during original sampling *ca.* 1970 are presented in Tables S2.3 & S2.4, which help in interpreting the observed rates of change.

We used the R package *vegan* (version 2.5-6, Oksanen et al., 2019) to calculate temporal turnover in community composition using the version of Jaccard distance that takes into account species abundances. Plot-scale turnover was logit-transformed before analyses to satisfy the assumption of homoskedasticity. We also calculated temporal turnover for the entire dataset.

### Modelling

We used those sites for which we had the full set of both response variables (species richness, species diversity, species evenness, height, LDMC and SLA) and predictor variables (canopy cover and canopy SLA, reindeer densities, management intensity and climate). The final dataset thus consisted of 245 resampled sites. These sites were then assigned to five biogeographical districts (BGD, Fig. 1) based on the Finnish biogeographical province division to account for spatially structured confounders in modelling. The original provinces were slightly modified for this study by merging districts with very few plots to neighboring districts, and by moving plots between districts so as to make each district either completely within or outside the reindeer herding area.

We used hierarchical Bayesian regression models to analyze plot-level changes in species diversity and composition. Two kinds of models were deployed based on the nature of the response variable. For species richness, evenness, diversity, and the three functional trait composition variables, the response variables could be modelled as a function of time, whereas temporal turnover could only be modelled with spatially varying effects, because turnover already represents the difference between two timepoints. We built four models with increasing complexity to answer our stated questions. Model 1 investigates how diversity and composition have changed in the sampled forests on average. Model 2 studies if there is spatial variation in these vegetation changes. Model 3 tests if temporal trends in vegetation are related to changes in land-use, climate, and canopies. Finally, Model 4 studies if changes in species diversity are related to changes in functional composition. Because there were no average changes in species richness, and because species diversity is the product of richness and evenness, trends in diversity could not be strongly driven by changes in richness. Model 4 was thus only fitted for evenness. As average species size is directly related to the productivity and physical structure of the community, the model includes average species height as the only predictor describing functional composition. All responses were standardized by subtracting the mean and dividing by standard deviation before analyses. The structure for the expectation function of different models is presented in Table 1. Note that responses (with the exception of turnover) were fed to the model separately for each time period, i.e changes in diversity and functional composition were not pre-calculated. This was done to allow pooling of information across time and space, and to describe the data-generating process as accurately as feasible.

**Table 1:** Structure of the expectation function of models used in this paper. All models had a gaussian error distribution. Community property refers to standardized species richness, species diversity, species evenness, and the abundance-weighted averages of log-transformed SLA, LDMC and height. Turnover refers to standardized logit-transformed temporal Jaccard distance. See methods for descriptions of the different parameters (a,b,c,CLI,REI,CSLA,CCOV,MI,HEI). SLA = Specific leaf area, LDMC = Leaf dry matter content.

| Response | Model | Model structure |
| --- | --- | --- |
| E(Community property) | 1 | a[T] |
|  | 2 | Model 1 + b[BGD,t] + c[plot] |
| | 3 | Model 2 + CLI × Climate change + REI × Reindeer density change + CSLA × Canopy SLA change + CCOV[Canopy cover change] + MI[Management intensity class] |
| E(Evenness) | 4 | Model 3 + HEI × Height change |
| E(Turnover) | 1 | a |
|  | 2 | Model 1 + b[BGD] |
| | 3 | Model 2 + CLI × Climate change + REI × Reindeer density change + CSLA × Canopy SLA change + CCOV[Canopy cover change] + MI[Management intensity class] |

All models assumed a gaussian error distribution. In the models in Table 1, parameters b and c are BGD- and plot-level (random) effects, respectively. In models of composition and diversity, b pools information across space and time, so that all BGD-level effects within a time are shrunk towards a common mean, and BGD-level effects in different times correlate with each other. The parameter c pools information across to shrink plot-level effects towards a common mean. In models of turnover, b pools information across BGDs. CLI, REI, CSLA, CCOV, MI, and HEI are parameters that define the effects of explanatory variables on plant community properties.

To look at the total effects of forest management, we also built a model which left out changes in canopy cover and canopy SLA. Because the results of this model were very similar to those of Model 3, we only present results for Model 3.

In Models 3 and 4 of community properties, all environmental and land-use terms were set to zero for the original sample. Thus, the terms that are in model 3 but not in model 2 only affect composition in the resample. The continuous variables Climate change, Reindeer density change, Canopy SLA change and Height change were entered as a difference in the relevant quantity between resample and original time, and their parameter estimates thus describe changes in the community-level property if the given predictor increases by one standardized unit in time. Management intensity was entered as a categorical variable with 4 levels, and canopy cover change was entered as a factor with three levels: decrease, no change, increase. The effect of no canopy cover change was fixed to zero.

Bayesian hierarchical models were built using the R package for Bayesian modelling *rethinking* (version 2.11, McElreath, 2020), an interface to the probabilistic programming language Stan (Carpenter et al., 2017). In all models, coefficients for continuous variables were given a Normal(0,0.5) prior distribution, coefficients for categorical variables were given a Normal(0,1) prior distribution, all standard deviations were given an Exponential(1) prior distribution, and temporal correlations in the BGD-level random effect were given an LKJ(2) prior. There were no divergent transitions in any of the MCMC chains in any of our models, which would have indicated unreliable estimates of the parameter posterior distributions (Carpenter et al., 2017). In addition, all parameters converged to a stationary posterior distribution (split chain R-hat < 1.01).

To test whether vegetation height is related to evenness in time (modelled above) as well as in space, we modelled evenness as a function of height using a generalized additive model (Wood, 2011). The effect of height was entered as a thin plate spline with basis dimension 20. We also tested whether the height-evenness relationship was different between sampling times (whether there was a time-height interaction), but a comparison of AICs indicated this was not the case.

### Data availability

The data and scripts used to produce these results have been deposited to Zenodo (Happonen et al., 2021).

## Results

### Question 1: Overall vegetation changes

Examining average values across all biogeographical districts we detected no changes in species richness, but observed increases in plot-level Shannon diversity and evenness during the sampling interval. Average LDMC increased, but there was no detectable change in average species SLA or height (Fig. 2). Average plot-scale turnover between the sampling periods was about 71% (Fig. 2). For the full dataset (i.e. the relative abundances of all species across the study area), temporal Jaccard dissimilarity was 37%, which, when divided over the average sampling interval (44 years) using the equation for exponential decay (Long-term turnover = 1 - (1 - annual turnover)^44^), corresponded to an annual turnover of 2.2%.

**Fig. 2:**
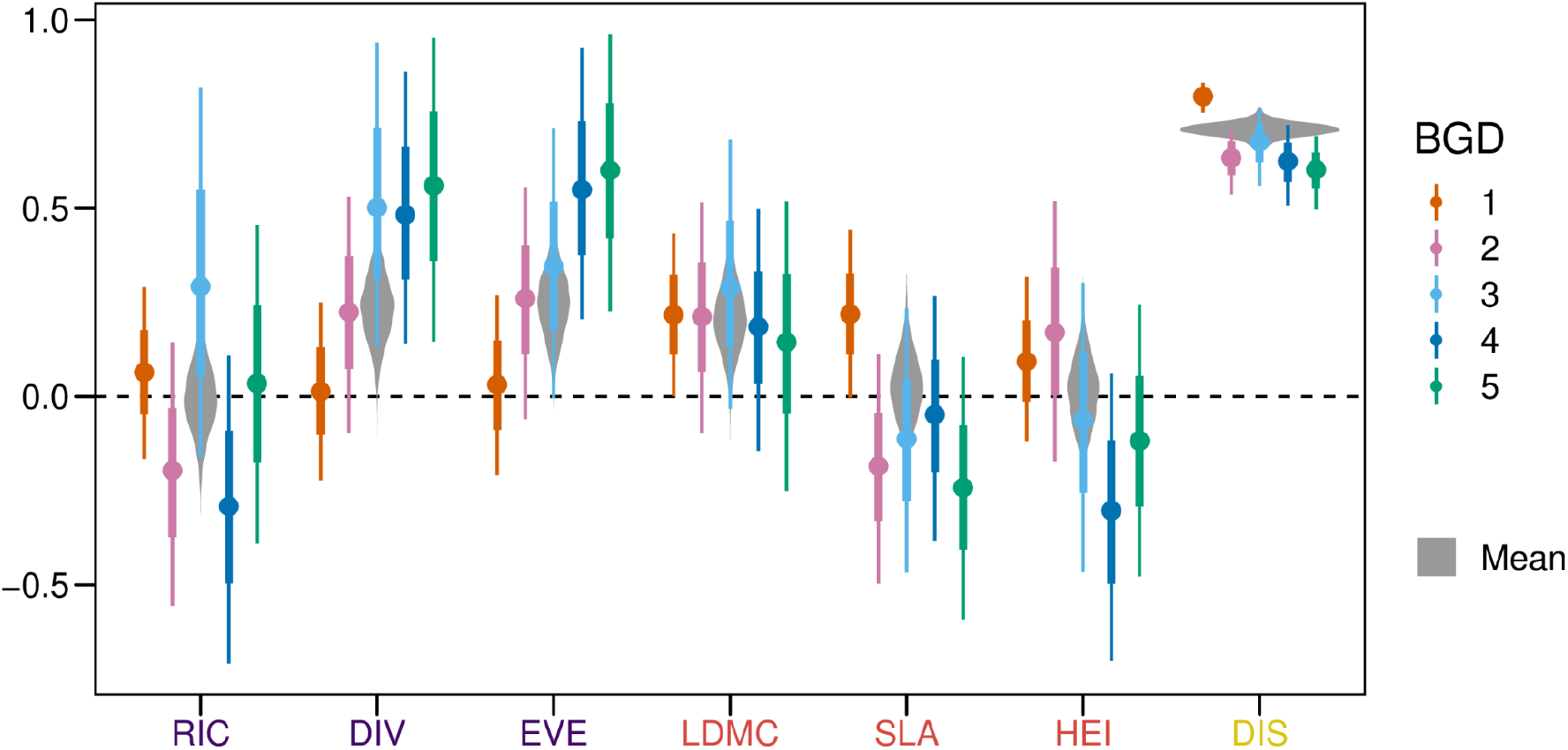
Temporal changes (1968–2019) in boreal herb-rich forest understory species richness (RIC), Shannon diversity (DIV), species evenness (EVE), lead dry matter content (LDMC), specific leaf area (SLA) and height (HEI), and average turnover (DIS), across the biogeographical districts (BGD), as inferred from model 2. The points are medians, the narrow and thick lines are 2/3 and 20/21 credible intervals, respectively, corresponding to 2/1 and 20/1 odds for the true parameter value to lie within the given interval, or 66.67% and 95.24% probability. The gray violin plot in the background is a kernel density plot of the posterior distribution of average change across all plots as inferred from model 1. All response variables except DIS are shown on a standardized scale. BGDs are arranged according to their latitude, from south (1) to north (5).

### Question 2: Geographical variation in vegetation changes

Vegetation changes across the study area were not representative of changes in all the BGDs (Fig. 2). Increases in species diversity and evenness had a clear geographic trend, with no changes in the south and clear increases in the north. Height changes also had a weak latitudinal gradient, with uncertain increases in height associated with more southern areas, and vice versa for the north. Patterns in species richness and SLA change and turnover were more idiosyncratic with respect to geography. Changes in LDMC did not differ geographically, but increased uniformly in all the BGDs.

### Question 3: Causes of vegetation changes

There was no discernible effect of observed climate change on any of the studied variables (Fig. 3a).

**Fig. 3:**
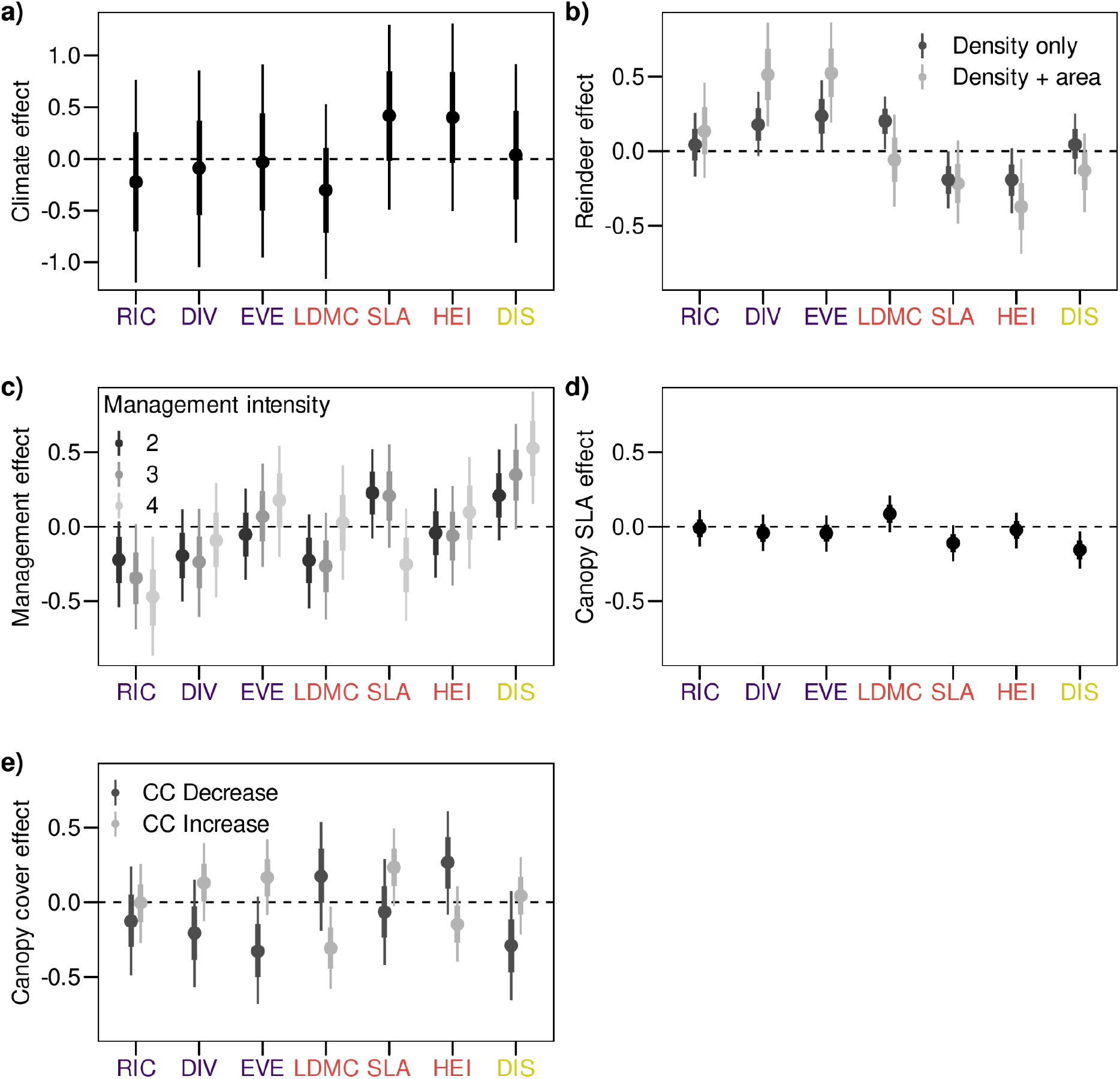
Standardized direct effects of climate change (a), reindeer (b), forest management intensity (c), canopy SLA increase (d), and canopy cover change (e) on community-level properties and turnover in boreal herb-rich forest understories, as inferred from model 3. Reindeer effects are displayed as the direct effect of reindeer density change and the combined effects of density change and being situated inside the reindeer herding area (average difference in trend compared to a plot outside the area). Management intensity effects are contrasts against unmanaged forests (intensity class 1). The points are medians, the narrow and thick lines are 2/3 and 20/21 credible intervals, respectively, corresponding to 2/1 and 20/1 odds for the true parameter value to lie within the given interval. Response abbreviations as in Fig. 2. Traits were log-transformed and turnover logit-transformed before analyses.

Increasing reindeer densities were associated with increasing species diversity, evenness and LDMC, and decreased SLA and height (Fig. 3b). In addition, plots within the reindeer herding area had more positive trends in diversity and evenness and more negative trends in height compared to those outside the reindeer herding area.

Forest management negatively affected species richness, and increased temporal turnover compared to unmanaged forests (Fig. 3c). The responses were reinforced by increasing management intensity. Temporal trends in trait composition were only weakly related to forest management. SLA showed a tendency to decrease and LDMC a tendency to decrease in the most intensively managed sites compared to natural sites.. Unmanaged forests had an uncertain positive richness trend, with a mean change of +0.3 standardized units and an 80% probability of the trend being positive.

Changes in canopy SLA correlated negatively with field-layer SLA and temporal turnover (Fig. 3d). Canopy SLA changes were almost completely explained by changes in the dominance of the evergreen tree *Picea abies* (Fig. S2.5), with *Picea* dominance corresponding to decreasing canopy SLA.

Canopy cover changes also affected these communities, with densening and sparsening canopies changing communities in different directions (Fig. 3e). Changes in canopy cover were positively correlated with changes in diversity, evenness, and SLA, and negatively with changes in LDMC and height.

### Question 4: Effect of height on diversity change

Height had a nonlinear but mostly negative spatial relationship with evenness (Fig. 4a). The temporal relationship between height and evenness was also negative (Fig. 4b). Including height as a predictor somewhat decreased the inferred effects of reindeer and canopy cover changes on evenness (Fig. S2.7).

**Fig. 4:**
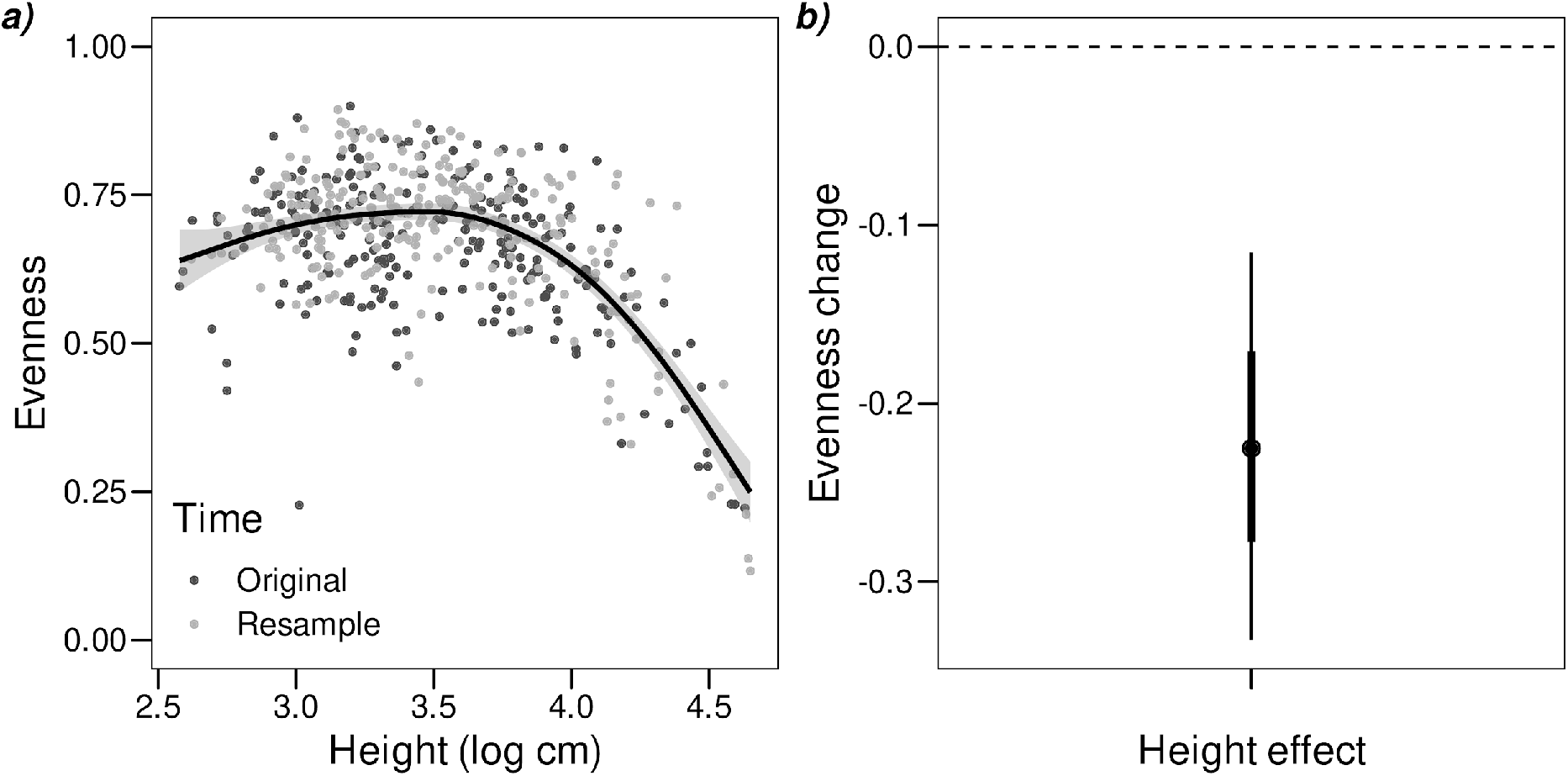
Spatial (a) and temporal (b) relationships between community evenness and average plant height. The spatial relationship is an unadjusted GAM of evenness against height. The temporal effect is a marginal standardized effect from hierarchical linear model 4 (see methods). In a), the line is the mean effect and the shading is the 20/21 confidence interval. In b), the point is the median effect, and the narrow and thick lines are 2/3 and 20/21 credible intervals, respectively. Refer to methods for more information.

## Discussion

We found that land-use, but not climate change, alters the functional composition of forest understory plant communities and explains part of the variation in temporal biodiversity trends both at regional and fine scales. Most evidently, increased reindeer densities had a clear effect on both average plant functional traits and diversity measures. Reindeer decreased average SLA and decreased plant height, which resulted in increased species evenness and diversity. Our finding shows that diversity changes were partially modulated by changes in plant size, illustrating a direct effect of changing functional composition on diversity. Our results thus highlight the way changes in average traits can be informative of changing selection pressures on vegetation, as most observed changes in functional composition were directly linked to land-use and canopy changes in ways that are consistent with theory. We thus provide evidence that including functional metrics in biodiversity monitoring programs is useful for inferring causes of biodiversity changes during the anthropocene (Pereira et al., 2013).

### Vegetation trends vary in space

Our analysis revealed that many community characteristics were stable over the 4-5 decades study interval across the study area, but displayed trends when vegetation changes were conditioned on geographic location. Diversity, evenness, and vegetation height had different trends in different parts of the latitudinal gradient. Apparent stability at larger scales can thus be the sum of opposing trends in different regions. We report that across the study area, species richness was on average stable as in a previous meta-analysis (Vellend et al., 2013), even though species turnover was on average very high (*ca.* 70%). Our results are thus consistent with the hypothesis that local species richness is highly constrained by regional processes such as compensatory colonization-extinction dynamics, meaning that lost species are readily replaced by new species from the metacommunity (Supp & Ernest, 2014). Average turnover within plots was high, most likely attributable to high species richness (Table S2.3), a long sampling interval, and inherent relocation error. Nonetheless, temporal turnover for the full dataset was of the same magnitude as average turnover in terrestrial ecosystems in a recent meta-analysis (2% year^−1^, Blowes et al., 2019), showing that the rate of compositional change in our data is representative of previously reported biodiversity changes.

### Land-use and canopy dynamics change the composition of field-layer vegetation

Our results provide clear evidence that reindeer control forest community composition by diminishing the relative abundance of tall plants with high SLA, thereby increasing species diversity, expanding the earlier findings from tundra and mountain forests (Kaarlejärvi, Eskelinen, & Olofsson, 2017; Sundqvist et al., 2019) to boreal forests. These findings are consistent with theoretical expectations, since SLA has been shown to correlate positively with nutrient concentrations in leaves (i.e. forage quality, Díaz et al., 2016), while large plants have been shown to be more susceptible to mammalian herbivory (Díaz et al., 2007). Furthermore, reindeer presence in itself seemed to affect vegetation changes even when animal densities remained the same, since changes in vegetation height and diversity were more pronounced in BGDs that were inside the reindeer herding area (BGDs 3, 4 and 5, Fig. 3). However, these effects could also be related to other confounding factors that change along the latitudinal gradient.

Surprisingly, forest management intensity did not have clear effects on functional composition. Instead, it decreased species richness and increased turnover. Forest management effects on turnover and diversity were mostly direct i.e. not mediated by management effects on canopy cover and composition, as evidenced by the low difference between management effects in models that included vs. excluded canopy variables. The finding of higher turnover in managed forests agrees with previous studies from boreal forests (Brice, Cazelles, Legendre, & Fortin, 2019; Kaarlejärvi, Salemaa, Tonteri, Merilä, & Laine, 2021) and other biomes (Barlow et al., 2016; Lake et al., 2000) reporting accelerated biodiversity change following human disturbance. Taken together, our results suggest that species richness changes in vascular plant communities in response to forest management do not seem to be driven by a filtering of species based on plant height or leaf traits. However, the highest management intensity class (class 4) that included recent clearcuts had more negative SLA and more positive LDMC trends compared to other managed forests, which supports previous findings of harvesting having a signal on the light-interception strategy of the understory community (Tonteri et al., 2016), but suggests that this effect is transient and undetectable in later forest development phases.

Canopy development also affected functional composition. Canopy cover had increased on average (see Fig.S2.6), leading to increased shading and decreased abundance of tall, light-demanding species with high SLA and low LDMC. SLA is a succession trait, which increases in response to shade (Dahlgren et al., 2006), whereas LDMC has probably decreased because of its negative covariance with SLA (Bruelheide et al., 2018). Canopy composition, expressed as canopy SLA, also influenced the functional composition of the understory. We would have expected changes in canopy SLA to correlate positively with nutrient availability and thus also with fast leaf economic traits (i.e. high SLA, low LDMC). Instead, we found the opposite. In our data, canopy SLA was almost perfectly negatively correlated with the relative abundance of *Picea abies* (see Fig.S2.5), which is the only abundant evergreen tree in these systems and frequently forms monodominant stands. This lack of diversity at one end of the trait spectrum limits the usefulness of functional traits in inferring the mechanisms by which canopy composition affects understories, since average trait metrics become perfectly correlated with all qualities of *Picea*. We hypothesize that in our data, the true mechanism behind the connection of canopy SLA and functional composition is increased shading with *Picea* dominance. *Picea* is evergreen and very shade tolerant, its canopy extends further down than that of other large trees, and it casts shade also during spring and autumn, when deciduous species are without leaves. This hypothesis is supported by the similar functional footprint of canopy SLA and shading on understories (decreased SLA, increased LDMC). The fact that decreased canopy SLA, through increased *Picea* dominance, was associated with increased community turnover agrees with previous literature that lists “borealization” as one threat against understories of semi-open, mixed or deciduous canopies (Kouki et al., 2018). Moreover, our results support the findings of a recent resurvey study from European temperate forests, which listed increasing canopy shade as the primary cause of changes in functional composition of herb-layers (Depauw et al., 2020), thereby generalizing these results across the temperate-boreal boundary.

### Changes in diversity are partially mediated by vegetation height

Changes in species diversity were mostly caused by changes in evenness, and those in turn were directly related to changes in vegetation height and its environmental drivers, namely reindeer herbivory and changes in canopy cover. Furthermore, the relationship between evenness and plant height was consistent both in space and time, providing further evidence that changes in functional composition can be a direct mechanism behind diversity changes. Reindeer grazing and canopy cover effects are most likely a result of the mechanistic link between size and diversity: tall species are being eaten or suppressed by lack of light, which reduces dominance, which in turn increases diversity (Grime, 1973; Oksanen, 1996). In these forests, both canopy cover and reindeer densities have increased on average. According to our results, both these changes favor shorter plants and result in increased evenness and diversity, thus partially explaining the observed large-scale changes in diversity. These results agree with Depauw et al. (2020), who also found that increased shading is responsible for increased evenness in the herb-layers of temperate European forests.

The effects of reindeer and canopy variables on evenness were not fully mediated by vegetation size (Fig. S2.7), because reindeer still had effects on diversity after adjusting for changes in plant height (model 4). One explanation may be that our study does not account for intraspecific changes in height, which could happen in response to altered light availability or grazing pressure (Jessen, Kaarlejärvi, Olofsson, & Eskelinen, 2020). Thus, the effects of reindeer herbivory and canopy cover on evenness via vegetation height are probably even stronger than detected here.

## Conclusions

Herbivory and canopy properties affect the diversity and composition of understory plant communities in boreal herb-rich forests over several decades (Fig. 5). Our results illustrate that land-use and canopy changes impact diversity indirectly by affecting average plant height. By favouring smaller species and limiting the size of plant individuals, increased shade and grazing pressure increase diversity. Therefore, plant size has a mechanistic connection with diversity. In addition, high light availability and reindeer densities favor slower leaf economic traits. Our results thus confirm that functional traits are both indicators and mediators of land-use change effects on plant communities, and underline the importance of including functional metrics in biodiversity monitoring programs.

**Fig. 5:**
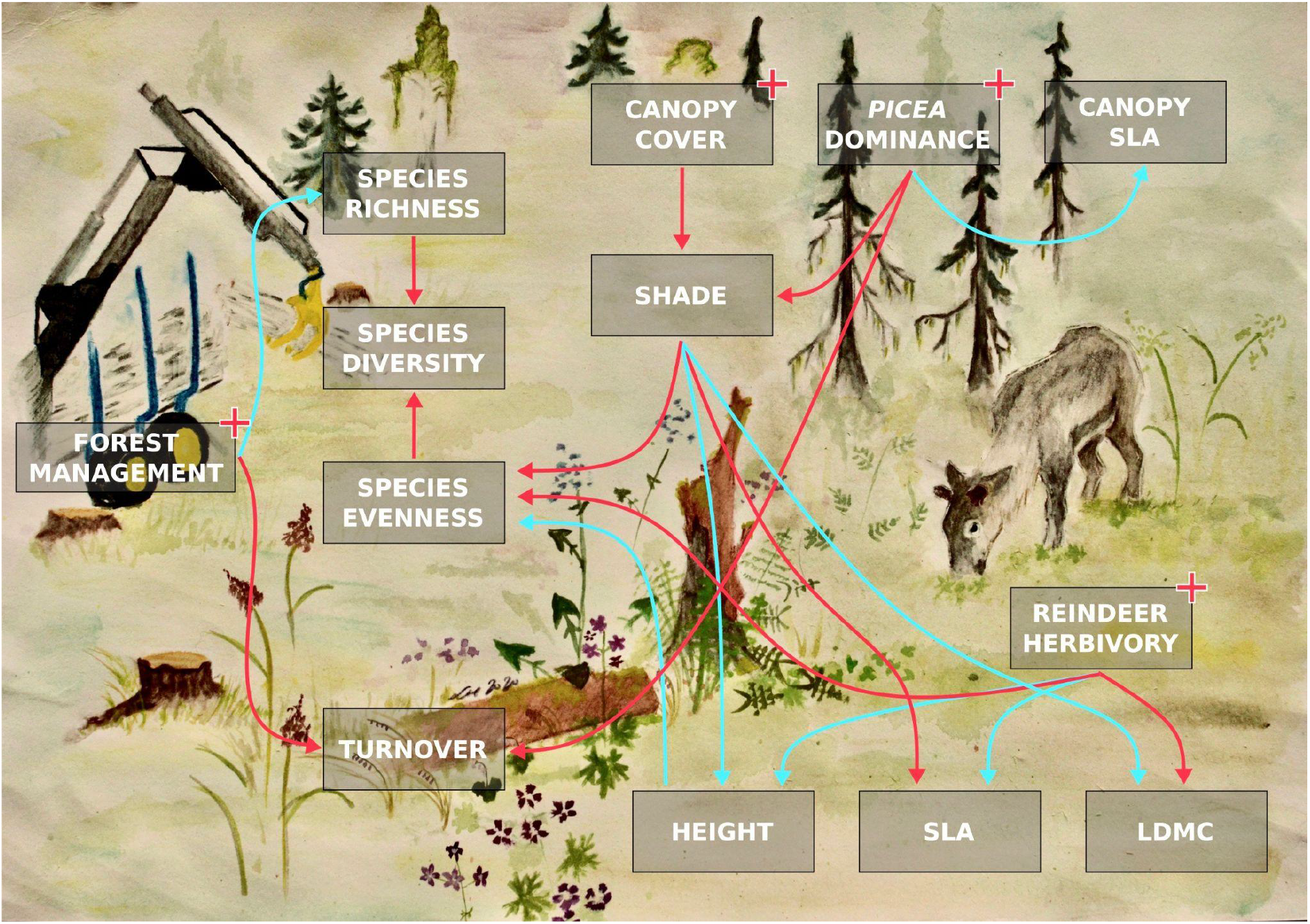
A conceptual model of vegetation change drivers in these boreal forests. Red lines correspond to positive effects, and blue lines to negative effects. Positive trends in land-use and canopy variables are marked with red plus signs. While based on our modelling results, the arrows are our subjective interpretation of their implications.

## Acknowledgments

We thank Jouko Kumpula from the Natural Resources Institute Finland for providing the reindeer density data. We acknowledge funding from Societas Pro Fauna et Flora Fennica, Societas Biologica Fennica Vanamo, Societas Amicorum Naturae Ouluensis, Academy of Finland (project #259072), Finnish Cultural Foundation, The Finnish Society of Forest Science and Jane & Aatos Erkko Foundation. We thank Anu Skog and Niilo Tenkanen for help in field work, and are grateful for the Botanical Museum and Oulanka Research station of the University of Oulu for providing facilities.

---

**Table S1.1:** Observed species and their trait values.

**Table S2.2:** The one-out-all-out criteria for inclusion in a management intensity class.

**Table S2.3:** Summary statistics for community properties during the original survey.

**Fig. S2.1:** Categorical predictors of vegetation change between 1968–1975 and 2013–2019.

**Fig. S2.2:** Distribution of changes in studied continuous drivers of vegetation change in standardized units.

**Fig. S2.3:** Harvests of large herbivores in Finland 2000–2018.

**Fig. S2.4:** Distributions of bioclimatic variables for the summer months in the study plots.

**Fig. S2.5:** The CWM of SLA in the tree layer is a function of the relative cover of Norway Spruce (*Picea abies*).

**Fig. S2.6:** Changes in the percentages of plots with high and low canopy cover, calculated by management intensity classes and biogeographical districts (BGD).

**Fig. S2.7:** All direct effects from Model 4.

## Notes

http://doi.org/10.5281/zenodo.4697167

### Competing Interest Statement

The authors have declared no competing interest.

### Summary of Updates

Text and figures reworked for clarity.

