## Appendix S1 for "Trait-based responses to land use and canopy dynamics modify long-term diversity changes in forest understories"

| Species | TPL record ID | Height (cm) | SLA ($mm^2 mg^{-1}$) | LDMC | Relative abundance |
|---|---|---|---|---|---|
| Achillea millefolium | gcc-140712 | 10.0 | 13.0 | 0.2 | 0.1% |
| Actaea rubra | kew-2620570 | 30.0 | 23.2 | 0.2 | 0.2% |
| Actaea spicata | kew-2620577 | 30.0 | 42.6 | 0.2 | 0.3% |
| Agrostis sp. | NA | 40.0 | 29.5 | 0.3 | 0.2% |
| Alchemilla sp. | NA | 16.0 | 19.1 | 0.3 | 0.1% |
| Anemone nemorosa | kew-2638482 | 15.0 | 25.0 | 0.2 | 0.0% |
| Angelica archangelica | kew-2638995 | 57.5 | 21.3 | 0.3 | 0.1% |
| Angelica sylvestris | kew-2639281 | 55.0 | 30.0 | 0.2 | 0.6% |
| Antennaria dioica | gcc-134033 | 1.5 | 23.6 | 0.2 | 0.0% |
| Anthoxanthum odoratum | kew-394021 | 14.5 | 26.7 | 0.3 | 0.0% |
| Anthriscus sylvestris | kew-2641868 | 60.0 | 40.8 | 0.2 | 0.5% |
| Athyrium filix-femina | tro-26600065 | 90.0 | 46.2 | 0.3 | 6.4% |
| Persicaria vivipara | kew-2572250 | 6.0 | 14.2 | 0.2 | 0.0% |
| Botrychium lunaria | tro-26602178 | 3.0 | 17.5 | 0.2 | 0.0% |
| Botrychium virginianum | tro-26602189 | 45.0 | NA | NA | 0.0% |
| Calypso bulbosa | kew-32237 | 1.0 | NA | NA | 0.0% |
| Calamagrostis arundinacea | kew-402396 | 90.0 | 17.7 | 0.4 | 0.0% |
| Calamagrostis canescens | kew-402519 | 120.0 | 21.6 | 0.4 | 0.1% |
| Calamagrostis epigejos | kew-402643 | 60.0 | 18.7 | 0.4 | 0.1% |
| Calamagrostis lapponica | kew-402934 | 15.0 | 25.7 | 0.4 | 0.0% |
| Calamagrostis purpurea | kew-403265 | 115.0 | 24.0 | 0.3 | 1.3% |
| Caltha palustris | kew-2693856 | 30.0 | 30.9 | 0.2 | 0.1% |
| Campanula rotundifolia | kew-365918 | 16.0 | 20.7 | 0.3 | 0.0% |
| Carex brunnescens | kew-225178 | 27.5 | 34.5 | 0.4 | 0.0% |
| Carex canescens | kew-225264 | 47.5 | 31.6 | 0.2 | 0.0% |
| Carex capillaris | kew-225305 | 15.0 | 13.2 | 0.4 | 0.0% |
| Carex cespitosa | kew-225431 | 42.5 | 25.8 | 0.3 | 0.1% |
| Carex digitata | kew-225999 | 10.0 | 27.4 | 0.2 | 0.3% |
| Carex disperma | kew-226066 | 61.0 | NA | NA | 0.0% |
| Carex echinata | kew-226238 | 27.5 | 15.0 | 0.4 | 0.0% |
| Carex sp. | NA | NA | NA | NA | 0.0% |
| Carex flava | kew-226588 | 40.0 | 22.0 | 0.2 | 0.0% |
| Carex globularis | kew-226981 | 37.5 | 18.8 | 0.4 | 0.0% |
| Carex loliacea | kew-228237 | 9.2 | NA | NA | 0.0% |
| Carex nigra | kew-229004 | 7.0 | 23.8 | 0.4 | 0.0% |
| Carex norvegica | kew-229087 | 13.5 | 40.9 | 0.3 | 0.0% |
| Carex leporina | kew-228079 | 30.0 | 22.4 | 0.3 | 0.0% |
| Carex pallescens | kew-229422 | 57.6 | 25.0 | 0.4 | 0.1% |
| Carex vaginata | kew-231861 | 22.5 | 19.3 | 0.3 | 0.2% |
| Carex vesicaria | kew-231937 | 80.0 | 16.5 | 0.3 | 0.0% |
| Centaurea phrygia | gcc-1229 | 55.0 | 30.5 | 0.1 | 0.0% |
| Cerastium fontanum | kew-2710319 | 25.0 | 30.5 | 0.2 | 0.0% |
| Lactuca alpina | gcc-100549 | 80.0 | 41.0 | 0.1 | 1.8% |
| Circaea alpina | kew-2721778 | 12.5 | 44.2 | 0.2 | 0.0% |
| Cirsium helenioides | gcc-104894 | 61.0 | 26.4 | 0.1 | 0.9% |
| Cirsium palustre | gcc-14233 | 92.5 | 20.1 | NA | 0.0% |
| Dactylorhiza viridis | kew-55635 | 15.0 | 21.8 | 0.2 | 0.0% |
| Convallaria majalis | kew-303076 | 22.2 | 34.3 | 0.2 | 1.6% |
| Cornus suecica | kew-47524 | 8.0 | 30.9 | 0.2 | 0.0% |
| Corallorhiza trifida | kew-47090 | 16.0 | NA | NA | 0.0% |
| Crepis paludosa | gcc-90103 | 62.5 | 37.0 | 0.1 | 1.2% |
| Cystopteris fragilis | tro-26600121 | 24.0 | 39.9 | 0.2 | 0.0% |
| Cystopteris montana | tro-26602776 | 25.0 | NA | NA | 0.0% |
| Dactylorhiza maculata | kew-55410 | 26.2 | 22.0 | 0.1 | 0.0% |
| Deschampsia cespitosa | kew-407579 | 81.0 | 13.9 | 0.3 | 0.6% |
| Deschampsia flexuosa | kew-407693 | 12.5 | 23.4 | 0.3 | 0.8% |
| Diplazium sibiricum | tro-26610748 | 41.5 | NA | NA | 0.1% |
| Dryopteris carthusiana | tro-26600144 | 50.0 | 38.8 | 0.2 | 0.4% |
| Dryopteris expansa | tro-26600148 | 80.0 | 44.0 | 0.3 | 3.6% |
| Dryopteris filix-mas | tro-26602214 | 120.0 | 24.9 | 0.3 | 0.0% |
| Elymus caninus | kew-410994 | 85.0 | 27.9 | 0.3 | 1.5% |
| Elymus repens | kew-411505 | 57.6 | 21.2 | 0.3 | 0.0% |
| Empetrum nigrum | kew-2788473 | 30.0 | 8.1 | 0.4 | 0.0% |
| Epilobium angustifolium | kew-2790112 | 100.0 | 14.2 | 0.2 | 0.5% |
| Epipogium aphyllum | kew-70443 | 10.0 | NA | NA | 0.0% |
| Epilobium hornemannii | kew-2790536 | 5.0 | NA | NA | 0.0% |
| Epilobium montanum | kew-2790730 | 45.0 | 26.7 | 0.2 | 0.0% |
| Epilobium palustre | kew-2790841 | 32.5 | 49.7 | 0.1 | 0.0% |
| Equisetum arvense | tro-26602003 | 42.4 | 12.1 | 0.2 | 0.1% |
| Equisetum fluviatile | tro-26602005 | 95.7 | 9.3 | 0.3 | 0.0% |
| Equisetum palustre | tro-26602010 | 35.0 | 8.0 | 0.2 | 0.0% |
| Equisetum pratense | tro-26602011 | 41.2 | 12.5 | 0.2 | 1.5% |
| Equisetum scirpoides | tro-26602012 | 12.5 | 9.3 | 0.3 | 0.0% |
| Equisetum sylvaticum | tro-26602013 | 27.4 | 24.4 | 0.3 | 1.6% |
| Eriophorum angustifolium | kew-244512 | 17.0 | 8.7 | 0.3 | 0.0% |
| Euphrasia frigida | tro-29204626 | 5.0 | 20.5 | NA | 0.0% |
| Festuca ovina | kew-415853 | 30.0 | 13.8 | 0.3 | 0.0% |
| Festuca rubra | kew-416502 | 50.0 | 12.1 | 0.3 | 0.0% |
| Filipendula ulmaria | rjp-1024 | 97.0 | 28.3 | 0.4 | 9.4% |
| Fragaria vesca | rjp-23 | 10.0 | 27.2 | 0.3 | 0.6% |
| Galium boreale | kew-85836 | 37.5 | 25.0 | 0.2 | 0.0% |
| Galeopsis bifida | kew-85339 | 47.5 | NA | NA | 0.0% |
| Galium trifidum | kew-87745 | 24.4 | NA | NA | 0.0% |
| Galium palustre | kew-87072 | 40.5 | 25.6 | 0.2 | 0.0% |
| Galium triflorum | kew-87764 | NA | 35.7 | 0.2 | 0.1% |
| Galium uliginosum | kew-87787 | 32.5 | 27.2 | 0.3 | 0.0% |
| Geranium sylvaticum | kew-2824138 | 30.0 | 21.7 | 0.2 | 13.7% |
| Geum rivale | rjp-357 | 37.5 | 18.8 | 0.2 | 0.8% |
| Glechoma hederacea | kew-90109 | 30.0 | 39.0 | 0.2 | 0.0% |
| Gnaphalium norvegicum | gcc-94096 | 8.0 | 20.4 | 0.2 | 0.0% |
| Goodyera repens | kew-91902 | 10.0 | 18.2 | 0.2 | 0.0% |
| Gymnocarpium dryopteris | tro-26600166 | 18.5 | 45.8 | 0.2 | 8% |
| Hepatica nobilis | tro-27100409 | 8.0 | 17.8 | 0.2 | 0.0% |
| Heracleum sphondylium | kew-2846164 | 100.0 | 19.6 | 0.2 | 0.0% |
| Hieracium sp. | NA | 32.5 | 26.6 | 0.1 | 0.2% |
| Humulus lupulus | kew-2855039 | 400.0 | 27.0 | 0.3 | 0.0% |
| Huperzia selago | tro-26603630 | 10.0 | 13.8 | 0.3 | 0.0% |
| Juncus filiformis | kew-315000 | 30.0 | 10.1 | 0.3 | 0.0% |
| Lathyrus pratensis | ild-7773 | 50.0 | 29.5 | 0.3 | 0.0% |
| Scorzoneroides autumnalis | gcc-121516 | 7.5 | 26.3 | 0.2 | 0.0% |
| Leucanthemum vulgare | gcc-135712 | 40.0 | 16.2 | 0.2 | 0.0% |
| Linnaea borealis | kew-2337127 | 5.0 | 25.4 | 0.3 | 0.5% |
| Neottia cordata | kew-134197 | 6.0 | 31.7 | 0.1 | 0.0% |
| Neottia ovata | kew-134271 | 45.0 | 23.1 | 0.1 | 0.0% |
| Luzula multiflora | kew-316484 | 15.0 | 29.3 | 0.2 | 0.0% |
| Luzula pilosa | kew-316622 | 20.0 | 25.9 | 0.2 | 0.3% |
| Luzula sudetica | kew-316741 | 15.0 | 32.5 | 0.2 | 0.0% |
| Lycopodium annotinum | tro-26600190 | 10.0 | 17.4 | 0.4 | 0.2% |
| Lysimachia thyrsiflora | kew-2492247 | 45.0 | 38.1 | 0.2 | 0.0% |
| Lysimachia vulgaris | kew-2492070 | 98.0 | 20.9 | 0.2 | 0.0% |
| Maianthemum bifolium | kew-281331 | 16.2 | 32.9 | 0.2 | 3% |
| Matteuccia struthiopteris | tro-26602315 | 110.0 | 27.2 | 0.2 | 4.8% |
| Melica nutans | kew-423708 | 35.0 | 9.8 | 0.3 | 2.7% |
| Melampyrum pratense | tro-29200344 | 30.0 | 26.1 | 0.2 | 0.0% |
| Melampyrum sylvaticum | kew-2507238 | 20.0 | 31.8 | 0.2 | 0.4% |
| Milium effusum | kew-424434 | 80.0 | 37.0 | 0.2 | 1.8% |
| Molinia caerulea | kew-424761 | 85.0 | 18.3 | 0.3 | 0.0% |
| Moneses uniflora | kew-2372127 | 7.5 | 16.2 | 0.3 | 0.1% |
| Myosotis sylvatica | kew-2358257 | 23.8 | 31.7 | 0.2 | 0.0% |
| Orthilia secunda | kew-2387708 | 12.0 | 18.0 | 0.3 | 0.4% |
| Oxalis acetosella | kew-2394737 | 8.5 | 60.7 | 0.1 | 3.8% |
| Parnassia palustris | kew-2559858 | 10.0 | 20.3 | 0.1 | 0.0% |
| Paris quadrifolia | kew-283886 | 25.0 | 35.0 | 0.1 | 1.4% |
| Phalaris arundinacea | kew-433393 | 146.9 | 19.4 | 0.3 | 0.0% |
| Phegopteris connectilis | tro-26600261 | 22.5 | 42.2 | 0.2 | 3.5% |
| Phleum alpinum | kew-433718 | 10.0 | 16.1 | 0.4 | 0.0% |
| Phleum pratense | kew-433846 | 61.2 | 20.4 | 0.3 | 0.0% |
| Platanthera bifolia | kew-157039 | 25.0 | 19.2 | 0.1 | 0.0% |
| Plantago major | kew-2569743 | 15.0 | 18.9 | 0.2 | 0.0% |
| Poa annua | kew-435194 | 15.0 | 36.7 | 0.2 | 0.0% |
| Poa nemoralis | kew-436600 | 45.3 | 32.1 | 0.3 | 0.1% |
| Poa pratensis | kew-436906 | 13.0 | 17.6 | 0.3 | 0.1% |
| Polygonatum odoratum | kew-284008 | 20.5 | 33.5 | 0.2 | 0.0% |
| Potentilla erecta | rjp-1192 | 21.8 | 24.4 | 0.3 | 0.0% |
| Comarum palustre | rjp-1293 | 30.5 | 15.6 | 0.3 | 0.0% |
| Prunella vulgaris | kew-166020 | 19.0 | 27.1 | 0.2 | 0.1% |
| Pteridium aquilinum | tro-26600295 | 70.0 | 11.1 | 0.2 | 0.2% |
| Pyrola media | kew-2410546 | 5.3 | 20.4 | 0.3 | 0.0% |
| Pyrola minor | kew-2410543 | 6.5 | 19.3 | 0.3 | 0.1% |
| Pyrola rotundifolia | kew-2410818 | 10.0 | 20.4 | 0.3 | 0.2% |
| Ranunculus acris | kew-2524461 | 9.0 | 21.1 | 0.2 | 0.2% |
| Ranunculus auricomus | kew-2524362 | 27.0 | 16.2 | 0.2 | 0.0% |
| Coptidium lapponicum | kew-2736094 | NA | NA | NA | 0.0% |
| Ranunculus repens | kew-2526688 | 20.0 | 23.1 | 0.2 | 0.1% |
| Rhinanthus minor | kew-2900773 | 30.0 | 15.2 | 0.2 | 0.0% |
| Rorippa palustris | kew-2417173 | 36.8 | 40.8 | 0.1 | 0.0% |
| Rubus arcticus | rjp-827 | 30.5 | 49.2 | 0.3 | 0.3% |
| Rubus chamaemorus | rjp-6 | 6.0 | 12.5 | 0.3 | 0.0% |
| Rubus saxatilis | rjp-7 | 15.0 | 32.9 | 0.2 | 6.4% |
| Rumex acetosa | kew-2424198 | 7.0 | 26.3 | 0.1 | 0.0% |
| Rumex acetosella | kew-2424185 | 10.0 | 19.5 | 0.1 | 0.0% |
| Saussurea alpina | gcc-119529 | 9.5 | 17.4 | 0.2 | 0.0% |
| Scutellaria galericulata | kew-189114 | 25.0 | 42.8 | 0.2 | 0.0% |
| Selaginella selaginoides | tro-26613766 | 9.0 | 23.3 | NA | 0.0% |
| Silene dioica | kew-2488209 | 27.5 | 48.3 | 0.1 | 0.3% |
| Solidago virgaurea | gcc-44550 | 10.0 | 31.1 | 0.2 | 1% |
| Stellaria borealis | tro-6301716 | NA | NA | NA | 0.0% |
| Stellaria graminea | kew-2482031 | 40.5 | 22.5 | 0.2 | 0.0% |
| Stellaria longifolia | kew-2481944 | 17.5 | NA | NA | 0.0% |
| Stellaria media | kew-2481938 | 30.0 | 37.5 | 0.1 | 0.0% |
| Stellaria nemorum | kew-2482625 | 40.0 | 42.7 | 0.1 | 0.4% |
| Stellaria sp. | NA | NA | NA | NA | 0.0% |
| Taraxacum sp. | NA | 18.3 | 24.3 | 0.2 | 0.0% |
| Thalictrum flavum | kew-2512664 | 65.0 | 26.1 | 0.2 | 0.0% |
| Lysimachia europaea | kew-2896575 | 11.2 | 25.0 | 0.2 | 1.4% |
| Trifolium pratense | ild-8127 | 33.0 | 24.9 | 0.3 | 0.0% |
| Trifolium repens | ild-8135 | 20.0 | 29.5 | 0.2 | 0.1% |
| Trollius europaeus | kew-2514617 | 32.5 | 25.6 | 0.2 | 0.3% |
| Tussilago farfara | gcc-78510 | 17.5 | 13.9 | 0.1 | 0.0% |
| Urtica dioica | kew-2448560 | 102.7 | 21.1 | 0.3 | 0.2% |
| Vaccinium myrtillus | kew-2457648 | 25.0 | 18.5 | 0.4 | 1.2% |
| Vaccinium uliginosum | kew-2457998 | 30.0 | 13.4 | 0.4 | 0.0% |
| Vaccinium vitis-idaea | kew-2457314 | 16.4 | 7.4 | 0.4 | 1.3% |
| Valeriana sambucifolia | tro-33500013 | 73.5 | 30.9 | 0.1 | 0.1% |
| Veronica longifolia | kew-2462752 | 55.9 | 15.7 | 0.3 | 0.0% |
| Veronica officinalis | kew-2463283 | 10.0 | 21.4 | 0.3 | 0.0% |
| Vicia cracca | ild-9103 | 75.0 | 22.7 | 0.3 | 0.0% |
| Vicia sepium | ild-7863 | 47.5 | 25.3 | 0.2 | 0.0% |
| Viola canina | kew-2463879 | 18.0 | 30.2 | 0.2 | 0.0% |
| Viola epipsila | kew-2460299 | 11.5 | 43.5 | 0.2 | 1.7% |
| Viola mirabilis | kew-2459423 | 20.0 | 45.1 | 0.2 | 0.3% |
| Viola palustris | kew-2459417 | 7.8 | 34.6 | 0.2 | 0.0% |
| Viola riviniana | kew-2459647 | 10.0 | 35.8 | 0.3 | 0.0% |
| Viola selkirkii | kew-2459696 | 7.8 | 9.0 | 0.1 | 0.2% |
