## Appendix S2 for "Trait-based responses to land use and canopy dynamics modify long-term diversity changes in forest understories"

### Appendix S2: supplementary material for the paper ‘Trait-based responses to forestry and reindeer husbandry modify long-term changes in forest understories’

#### Tables

| | Natural sites = 1 | Semi-natural sites = 2 | Managed sites = 3 | Heavily managed sites = 4 |
|---|---|---|---|---|
| MANAGEMENT INTENSITY | Not managed | Light management | Intensive management (but no ditching or mounding) | Very intensive management (incl. ditching, fertilization) |
| SOIL | Undisturbed | Disturbed, but enables prevalence of herb-rich forest species | Disturbed, declined conditions for herb-rich forest species prevalence | Severely disturbed, soil may have dried up or topsoil eroded, prevalence of herb-rich forest species endangered |
| VEGETATION LAYERS | All layers present | Layer variability declined | Layer variability clearly declined | Layers missing |
| STAND STRUCTURE | Natural variation, gap dynamic works | Variability declined, impacts of management visible in gap dynamics | Pronounced decline in variability | Even-aged stand, plantation |
| DEADWOOD | Varying aged and sized | Impacts of management visible in the amount of decaying wood | Scarce or invariable in age or size | Scarce or missing |

**Table S2.2:** The one-out-all-out criteria for inclusion in a management intensity (Kouki et al. 2018).

| Variable | Unit | Mean | SD | Min. | Max. |
|---|---|---|---|---|---|
| Species richness | Species | 21.97 | 5.99 | 8.00 | 44.00 |
| Species Diversity | Species | 8.24 | 3.49 | 1.77 | 28.89 |
| Evenness | Unitless | 0.66 | 0.12 | 0.22 | 0.90 |
| Height | Log cm | 3.50 | 0.45 | 2.58 | 4.63 |
| LDMC | Log proportion | -1.49 | 0.12 | -1.81 | -1.11 |
| SLA | Log $mm^2mg^{-1}$ | 3.38 | 0.18 | 2.88 | 3.82 |
| Turnover | Logit proportion | 1.43 | 0.89 | -2.28 | 4.02 |

**Table S2.3:** Summary statistics for community properties during the original survey (*ca.* 1970) and for turnover. LDMC: leaf dry matter content, SLA: specific leaf area.

### Figures

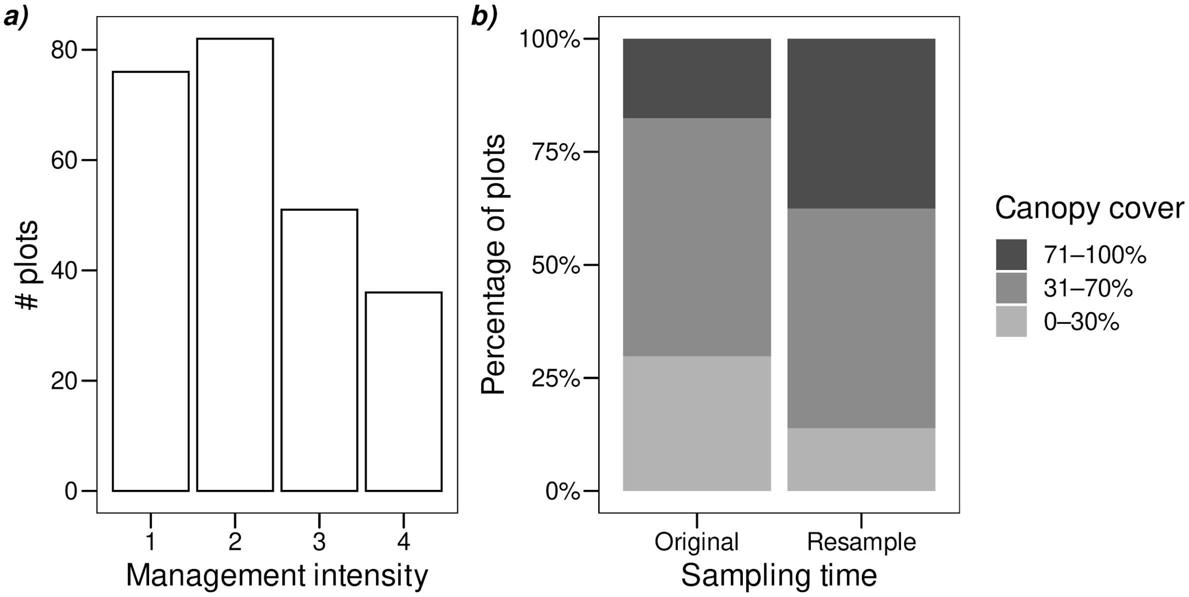


**Fig. S2.1:** Categorical predictors of vegetation change between 1968–1975 and 2013–2019, including the distribution of management intensity classes during the resurvey (a), and the distribution of canopy cover classes during both survey times (b).

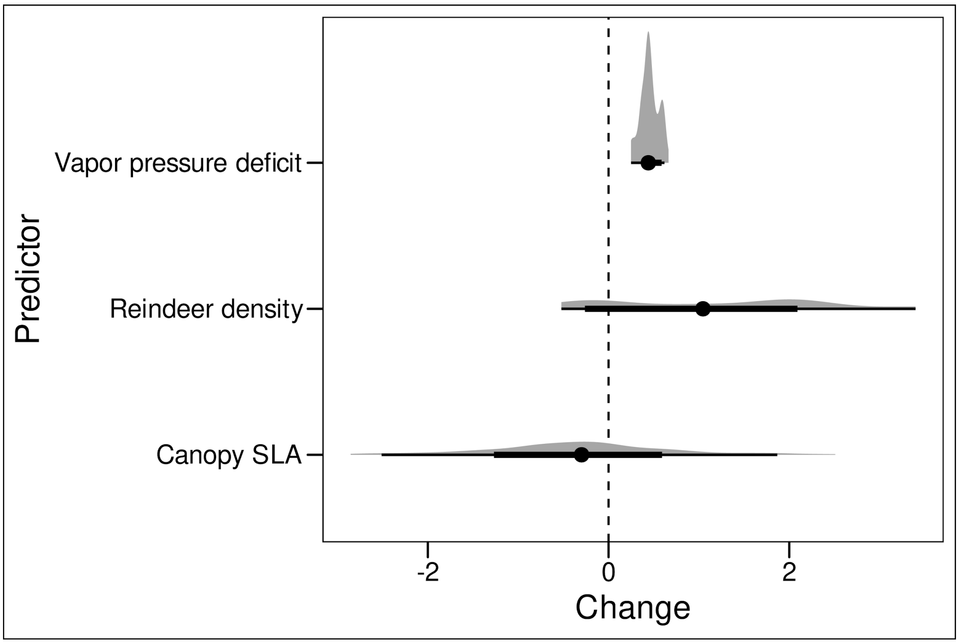


**Fig. S2.2:** Distribution of changes in studied continuous drivers of vegetation change in standardized units. The gray shading is a kernel density plot, the points are means, the thick (thin) lines depict ⅔ (20/21) quantile intervals, i.e. the change is 2 (20)  times more likely to be within the interval than outside it. Vapor pressure deficit change the difference between averages of years 1961–1970 and 2006–2015 from the Terra Climate dataset. Reindeer density change is based on yearly reindeer density data the Natural Resources Institute of Finland, and calculated by comparing 10-year averages prior to sampling of each plot. Canopy SLA is the change in abundance-weighted  average SLA on the log scale between sampling times. All values were standardized by the standard deviation of the variable during original sampling.

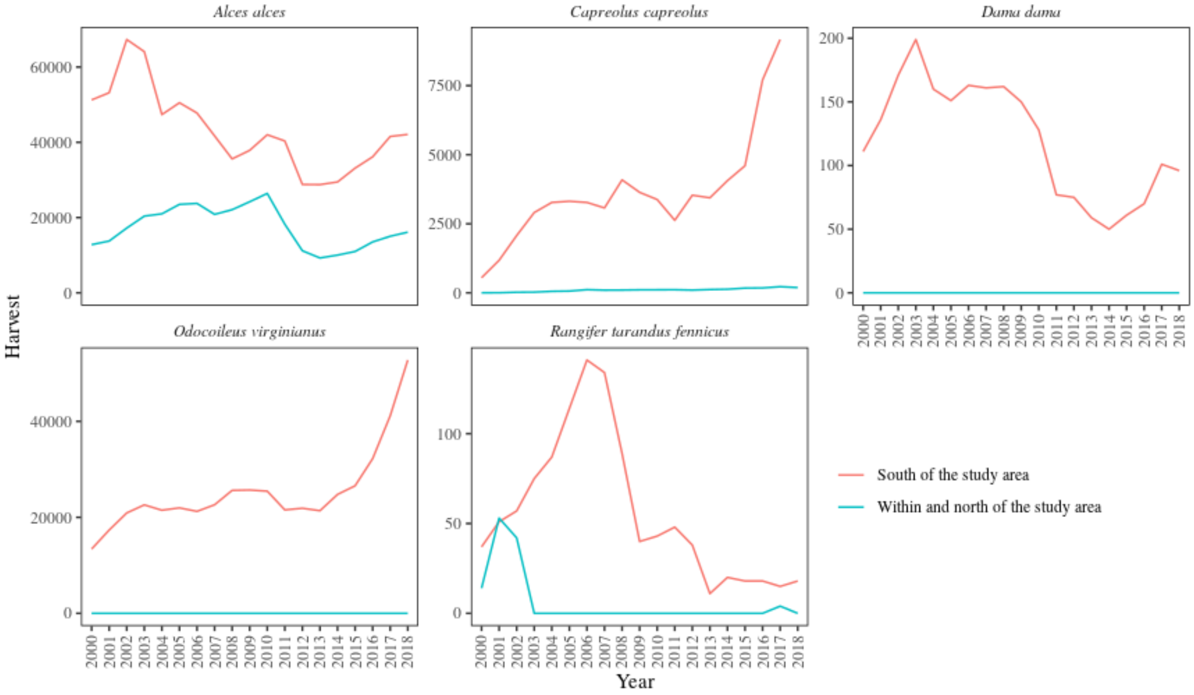


**Fig. S2.3:** Harvests of large herbivores in Finland 2000–2018, within and outside the study area. Data are from Natural Resources Institute Finland (2020).

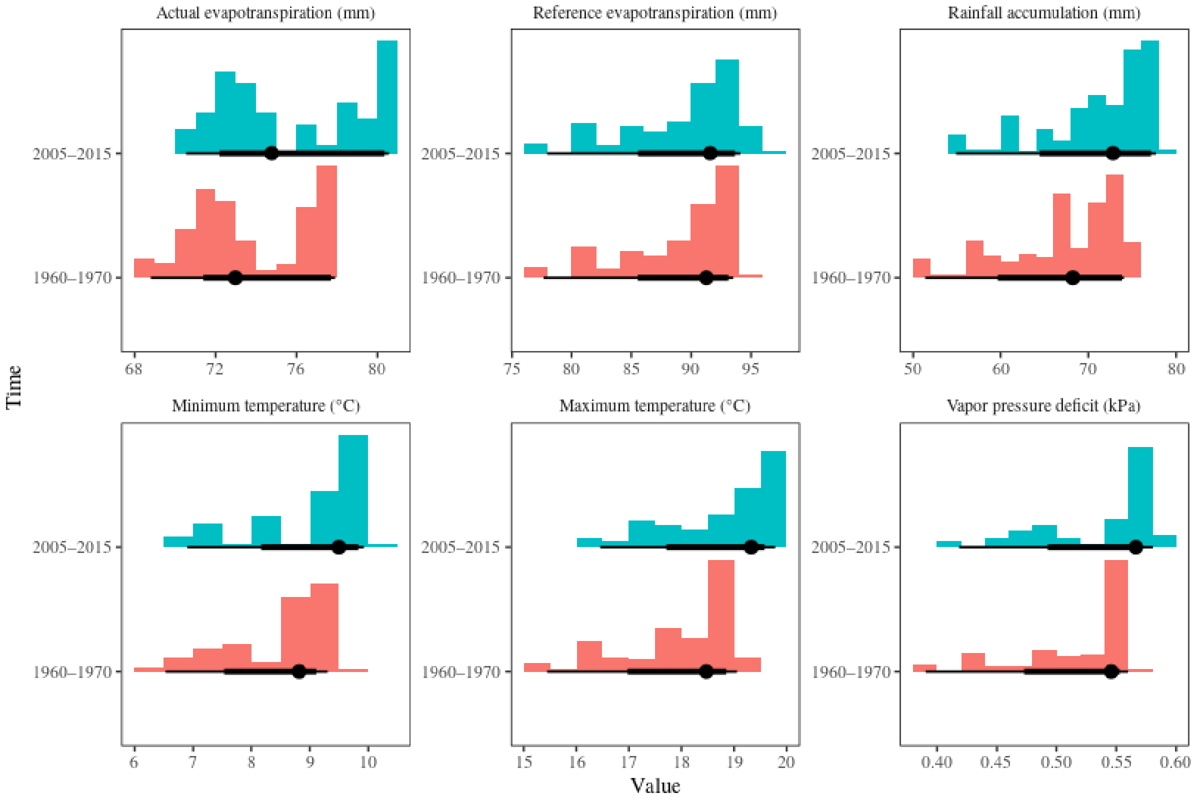


**Fig. S2.4:** Distributions of bioclimatic variables for the summer months in the study plots. The values are 10-year averages from the TerraClimate dataset (Abatzoglou et al., 2018), which has a nominal resolution of 1/24 degrees (~4km). The coloured bars are histograms, the points are medians, and the narrow and thick lines depict 95% and 50% quantile intervals, respectively.

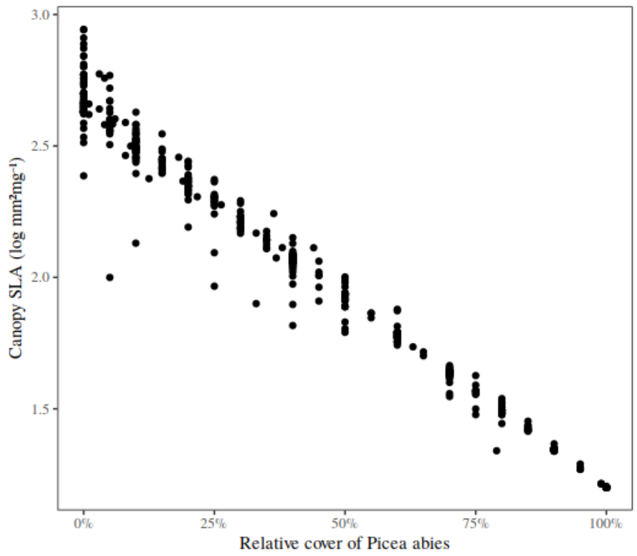


**Fig. S2.5:** The CWM of SLA in the tree layer is a function of the relative cover of Norway Spruce (*Picea abies*). SLA: specific leaf area.

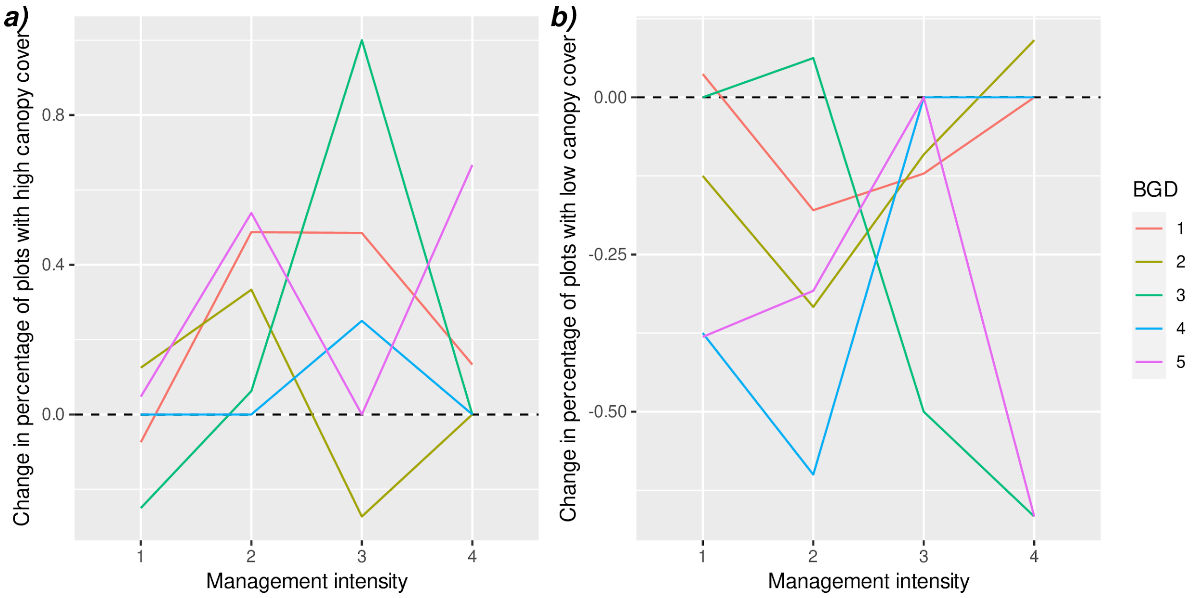


**Fig. S2.6:** Changes in the percentages of plots with high (a) and low (b) canopy cover, calculated by management intensity classes and biogeographical districts (BGD). There is a general trend toward higher canopy cover in all management intensity classes.

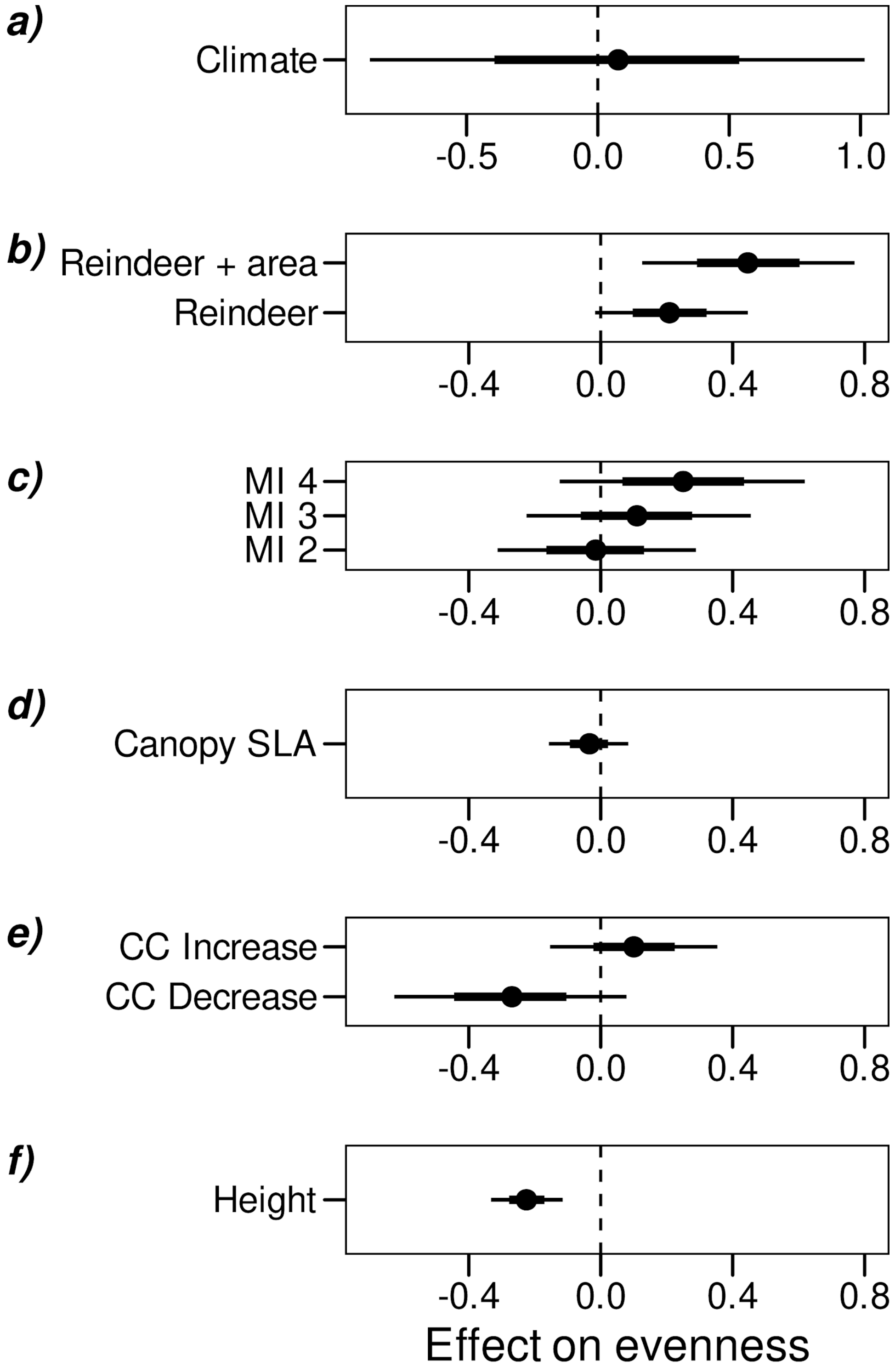


**Fig. S2.7:** All direct effects from Model 4, where evenness was the only response variable. MI = Management intensity, SLA = Specific leaf area, CC = canopy cover.
