## Appendix S3 for "Trait-based responses to land use and canopy dynamics modify long-term diversity changes in forest understories"

### Appendix S3: Original references for trait data from the TRY and LEDA databases used in the paper “Trait-based responses to forestry and animal husbandry modify long-term changes in forest understories”

References are as supplied in the databases.

Arrendondo, J. Tulio (2003): Components of leaf elongation rate and their relationship to specific leaf area in ...

Atkin, O. K., M. H. M. Westbeek, M. L. Cambridge, H. Lambers, and T. L. Pons. 1997. Leaf respiration in light and darkness - A comparison of slow- and fast-growing Poa species. Plant Physiology 113:961-965.

BIOPOP April 2005

Blonder, B., Buzzard, B., Sloat, L., Simova, I., Lipson, R., Boyle, B., Enquist, B. (2012) The shrinkage effect biases estimates of paleoclimate. American Journal of Botany. 99.11 1756-1763

Bond-Lamberty, B., C. Wang, and S. T. Gower (2002), Leaf area dynamics of a boreal black spruce fire chronosequence, Tree Physiol., 22(14), 993-1001.

Bragazza L (2009) Conservation priority of Italian alpine habitats: a floristic approach based on potential distribution of vascular plant species. Biodiversity and Conservation 18: 28232835.

Bruun, Hans Henrik(2005): Distinct patterns in alpine vegetation around dens of the Arctic fox [28]

Burrascano S, Copiz R, Del Vico E, Fagiani S, Giarrizzo E, Mei M, Mortelliti A, Sabatini FM, Blasi C (2015) Wild boar rooting intensity determines shifts in understorey composition and functional traits. COMMUNITY ECOLOGY 16(2) 244-253 DOI: 10.1556/168.2015.16.2.12

Campbell, C., L. Atkinson, J. Zaragoza-Castells, M. Lundmark, O. Atkin, and V. Hurry. 2007. Acclimation of photosynthesis and respiration is asynchronous in response to changes in temperature regardless of plant functional group. New Phytologist 176:375-389.

Campbell, Daniel R.(2003): Germination and seedling growth of bog plants in relation to the recolonization of milled peatlands [169]

Campetella, G; Botta-Dukát, Z; Wellstein, C; Canullo, R; Gatto, S; Chelli, S; Mucina, L; Bartha, S (2011): Patterns of plant trait-environment relationships along a forest succession chronosequence. Agriculture, Ecosystems & Environment, 145(1), 38-48. doi:10.1016/j.agee.2011.06.025

Ciocarlan V. (2009). The illustrated Flora of Romania. Pteridophyta et Spermatopyta. Editura Ceres, 1141 p (in Romanian).

Cody, W. J.(1975): The biology of canadian weeds. 15. Pteridium aquilinum (L.) Kuhn [55]

Cornelissen, J. H. C. 1996. An experimental comparison of leaf decomposition rates in a wide range of temperate plant species and types. Journal of Ecology 84:573-582.

Cornelissen, J. H. C., B. Cerabolini, P. Castro-Diez, P. Villar-Salvador, G. Montserrat-Marti, J. P. Puyravaud, M. Maestro, M. J. A. Werger, and R. Aerts. 2003. Functional traits of woody plants: correspondence of species rankings between field adults and laboratory-grown seedlings? Journal of Vegetation Science 14:311-322.

Cornelissen, J. H. C., H. M. Quested, D. Gwynn-Jones, R. S. P. Van Logtestijn, M. A. H. De Beus, A. Kondratchuk, T. V. Callaghan, and R. Aerts. 2004. Leaf digestibility and litter decomposability are related in a wide range of subarctic plant species and types. Functional Ecology 18:779-786.

Cornelissen, J. H. C., P. C. Diez, and R. Hunt. 1996. Seedling growth, allocation and leaf attributes in a wide range of woody plant species and types. Journal of Ecology 84:755-765.

Craine JM, Nippert JB, Towne EG, Tucker S, Kembel SW, Skibbe A, McLauchlan KK (2011) Functional consequences of climate-change induced plant species loss in a tallgrass prairie. Oecologia 165: 1109-1117

Craine, J.M.(201): The relationship among roots and leaf traits of 76 grassland species and relative abundance ...

Dale, M. P.(1992): The ecophysiology of Veronica chamaedrys, V. montana and V. officinalis. I. Light quality and light quantity

Dale, M. P.(1992): The ecophysiology of Veronica chamaedrys, Veronica montana and Veronica officinalis. II. The interaction of irradiance and water regime

Dalke, I.V., Novakovskiy, A.B., Maslova, S.P. et al. Plant Ecol (2018) 219: 1295. https://doi.org/10.1007/s11258-018-0879-2

De Vries F., Bardgett R.D. (2016) Plant community controls on short-term ecosystem nitrogen retention. New Phytologist. doi: 10.1111/nph.13832

den Dubbelden, Koen C.(1996): Inherent allocation patterns and potential growth rates of herbaceous climbing plants

Díaz, S., J. G. Hodgson, K. Thompson, M. Cabido, J. H. C. Cornelissen, A. Jalili, G. Montserrat-Martí, J. P. Grime, F. Zarrinkamar, Y. Asri, S. R. Band, S. Basconcelo, P. Castro-Díez, G. Funes, B. Hamzehee, M. Khoshnevi, N. Pérez-Harguindeguy, M. C. Pérez-Rontomé, F. A. Shirvany, F. Vendramini, S. Yazdani, R. Abbas-Azimi, A. Bogaard, S. Boustani, M. Charles, M. Dehghan, L. de Torres-Espuny, V. Falczuk, J. Guerrero-Campo, A. Hynd, G. Jones, E. Kowsary, F. Kazemi-Saeed, M. Maestro-Martínez, A. Romo-Díez, S. Shaw, B. Siavash, P. Villar-Salvador, and M. R. Zak. 2004. The plant traits that drive ecosystems: Evidence from three continents. Journal of Vegetation Science 15:295-304.

ECOFLORA - database of the ecological flora of the british isles

Eriksson, Ove(2003): Recruitment and life-history traits of sparse plant species in subalpine grasslands [81]

Everwand G, Fry, EL, Eggers T, Manning P (2014) Seasonal variation in the relationship between plant traits and grassland carbon and water fluxes. Ecosystems 17, 1095-1108

Fitter, A. H. and H. J. Peat 1994. The Ecological Flora Database. Journal of Ecology 82:415-425.

Freschet, G. T., J. H. C. Cornelissen, R. S. P. van Logtestijn, and R. Aerts. 2010. Evidence of the plant economics spectrum in a subarctic flora. Journal of Ecology 98:362-373.

Fry, E.L., Power, S.A. Manning, P. (2014) Trait based classification and manipulation of functional groups in biodiversity-ecosystem function experiments. Journal of Vegetation Science, 25, 248-261.

Garnier, E.(2001): Consistency of species ranking based on functional leaf traits

Green, W. 2009. USDA PLANTS Compilation, version 1, 09-02-02. (http://bricol.net/downloads/data/PLANTSdatabase/) NRCS: The PLANTS Database (http://plants.usda.gov, 1 Feb 2009). National Plant Data Center: Baton Rouge, LA 70874-74490 USA.

Gulias, Javier(2003): Relationship between Maximum Leaf Photosynthesis, Nitrogen Content and Specific Leaf Area in Balearic Endemic and Non-endemic Mediterranean Species [92]

Herz, K., Dietz, S., Haider, S., Jandt, U., Scheel, D. & Bruelheide, H. (2017): Drivers of intraspecific trait variation of grass and forb species in German meadows and pastures. Journal of Vegetation Science 28: 705716. Doi: 10.1111/jvs.12534. Herz, K., Dietz, S., Haider, S., Jandt, U., Scheel, D. & Bruelheide, H. (2017): Predicting individual plant performance in grasslands. Ecology and Evolution 7: 89588965. DOI: 10.1002/ece3.3393

HILL, M.O., PRESTON, C.D. & ROY, D.B. (2004) PLANTATT - attributes of British and Irish Plants: status, size, life history, geography and habitats. Huntingdon: Centre for Ecology and Hydrology.

Kattge, J., W. Knorr, T. Raddatz, and C. Wirth. 2009. Quantifying photosynthetic capacity and its relationship to leaf nitrogen content for global-scale terrestrial biosphere models. Global Change Biology 15:976-991.

Kichenin et al. 2013. Contrasting effects of plant inter- and intraspecific variation on community-level trait measures along an environmental gradient. Functional Ecology, in press.

Kleyer, M., R. M. Bekker, I. C. Knevel, J. P. Bakker, K. Thompson, M. Sonnenschein, P. Poschlod, J. M. van Groenendael, L. Klimes, J. Klimesova, S. Klotz, G. M. Rusch, Hermy, M. , D. Adriaens, G. Boedeltje, B. Bossuyt, A. Dannemann, P. Endels, L. Götzenberger, J. G. Hodgson, A.-K. Jackel, I. Kühn, D. Kunzmann, W. A. Ozinga, C. Römermann, M. Stadler, J. Schlegelmilch, H. J. Steendam, O. Tackenberg, B. Wilmann, J. H. C. Cornelissen, O. Eriksson, E. Garnier, and B. Peco. 2008. The LEDA Traitbase: a database of life-history traits of the Northwest European flora. Journal of Ecology 96:1266-1274.

Laughlin, D. C., J. J. Leppert, M. M. Moore, and C. H. Sieg. 2010. A multi-trait test of the leaf-height-seed plant strategy scheme with 133 species from a pine forest flora. Functional Ecology 24:493-501.

Lavergne, Sébastian(2003): Do rock endemic and widespread plant species differ under the Leaf-Height-Seed...

Louault, F., V. D. Pillar, J. Aufrere, E. Garnier, and J. F. Soussana. 2005. Plant traits and functional types in response to reduced disturbance in a semi-natural grassland. Journal of Vegetation Science 16:151-160.

Loveys, B. R., L. J. Atkinson, D. J. Sherlock, R. L. Roberts, A. H. Fitter, and O. K. Atkin. 2003. Thermal acclimation of leaf and root respiration: an investigation comparing inherently fast- and slow-growing plant species. Global Change Biology 9:895-910.

Lynn, D.E.(2003): Survival of Ranunculus repens L. (Creeping Buttercup) in an Amphibious Habitat [91]

Meziane, D. and B. Shipley. 1999. Interacting determinants of specific leaf area in 22 herbaceous species: effects of irradiance and nutrient availability. Plant Cell and Environment 22:447-459.

Meziane, D.(1999): Interacting determinats of specific leaf area in 22 herbaceous species: effects of irradiance and nutrient availability [22]

Meziane, Driss(2001): Direct and Indirect Relationships Between Specific Leaf Area, Leaf Nitrogen and Leaf Gas Exchange. Effects of Irradiance and Nutrient Supply [88]

Milla & Reich 2011 Annals of Botany 107: 455465, 2011.

Niinemets, Ülo(2003): Leaf structure vs. nutrient relationship vary with soil onditions in temperate shrubs and trees

Onoda Y, Wright IJ, Evans JR, Hikosaka K, Kitajima K, Niinemets Ü, Poorter H, Tosesns T, Westoby M. (2017) Physiological and structural tradeoffs underlying the leaf economics spectrum. New Phytologist

Ordonez, J. C., P. M. van Bodegom, J. P. M. Witte, R. P. Bartholomeus, J. R. van Hal, and R. Aerts. 2010. Plant Strategies in Relation to Resource Supply in Mesic to Wet Environments: Does Theory Mirror Nature? American Naturalist 175:225-239.

Parsons, A.N.(1994): Growth responses of four sub-Arctic dwarf shrubs to simulated environmental change

Paula, S., M. Arianoutsou, D. Kazanis, Ç. Tavsanoglu, F. Lloret, C. Buhk, F. Ojeda, B. Luna, J. M. Moreno, A. Rodrigo, J. M. Espelta, S. Palacio, B. Fernández-Santos, P. M. Fernandes, and J. G. Pausas. 2009. Fire-related traits for plant species of the Mediterranean Basin. Ecology 90:1420.

Peco B., de Pablos I., Traba J. , & Levassor C. (2005) The effect of grazing abandonment on species composition and functional traits: the case of dehesa Basic and Applied Ecology, 6(2): 175-183

Poorter, Hendrik(1998): Photosynthetic nitrogen-use efficiency of species that differ inherently in specific leaf area[116]

Poorter, Hendrik(1999): A comparison of specific leaf area, chemical composition and leaf construction costs of field ....

Prentice, I.C., Meng, T., Wang, H., Harrison, S.P., Ni, J., Wang, G., 2011. Evidence for a universal scaling relationship of leaf CO2 drawdown along a moisture gradient. New Phytologist 190: 169180

Price, C.A. and B.J. Enquist. Scaling of mass and morphology in Dicotyledonous leaves: an extension of the WBE model. (2007) Ecology 88(5): 11321141.

Pyankov, V. I., A. V. Kondratchuk, and B. Shipley. 1999. Leaf structure and specific leaf mass: the alpine desert plants of the Eastern Pamirs, Tadjikistan. New Phytologist 143:131-142.

Quested, H. M., J. H. C. Cornelissen, M. C. Press, T. V. Callaghan, R. Aerts, F. Trosien, P. Riemann, D. Gwynn-Jones, A. Kondratchuk, and S. E. Jonasson. 2003. Decomposition of sub-arctic plants with differing nitrogen economies: A functional role for hemiparasites. Ecology 84:3209-3221.

Reich, Peter B.(1998): Relationship of leaf dark respiration to leaf nitrogen, specific leaf area and leaf life-span:..

Sandel, B., J. D. Corbin, and M. Krupa. 2011. Using plant functional traits to guide restoration: a case study in California coastal grassland. Ecosphere 2(2):art23. doi:10.1890/ES10-00175.1

Schweingruber, F.H., Landolt, W.: The Xylem Database. Swiss Federal Research Institute WSL Updated (2005)

Sheremetev S.N. (2005) Herbs on the soil moisture gradient (water relations and the structural-functional organization). KMK, Moscow, 271 pp. (In Russian)

Shipley B., 2002. Trade-offs between net assimilation rate and specific leaf area in determining relative growth rate: relationship with daily irradiance, Functional Ecology(16) 682-689

Shipley, B. 1995. Structured Interspecific Determinants of Specific Leaf-Area in 34 Species of Herbaceous Angiosperms. Functional Ecology 9:312-319.

Shipley, B. and M. J. Lechowicz. 2000. The functional co-ordination of leaf morphology, nitrogen concentration, and gas exchange in 40 wetland species. Ecoscience 7:183-194.

Shipley, B. and T. T. Vu. 2002. Dry matter content as a measure of dry matter concentration in plants and their parts. New Phytologist 153:359-364.

Shipley, B.(1995): Structured interspecific determinants of specific leaf area in 34 species of herbaceous angiosperms [9]

Shipley, B.(2002): Trade-offs between net assimilation rate and specific leaf area in determining relative growth rate: relationship with daily irradiance[16]

Sophie Gachet, Errol Véla, Thierry Tatoni, 2005, BASECO: a floristic and ecological database of Mediterranean French flora. Biodiversity and Conservation 14(4):1023-1034

Source data from Carl von Ossietzky university of Oldenburg, Landscape Ecology Group, DE (Krüger), Correspondig address:

Source data from Carl von Ossietzky university of Oldenburg, Landscape Ecology Group, DE (Kühner), Corresponding address:

Source data from Norwegian Institute for Nature Research, NINA-Trondheim, NO (Aarnes), corresponding address:

Source data from University of Groningen, Community and Conservation Ecology Group, NL (Steendam), Corresponding address:

Source data from University of Regensburg, Chair of Botany, DE (Gesing), corresponding adress:

Spasojevic, M. J. and K. N. Suding. 2012. Inferring community assembly mechanisms from functional diversity patterns: the importance of multiple assembly processes. Journal of Ecology 100:652-661.

Storkey, J.(2004): Modelling Seedling Growth Rates of 18 Temperate Arable Weed Species as a Function of the Environment and Plant Traits [93]

Thuiller W - Traits of European Alpine Flora - Wilfried Thuiller - OriginAlps Project - Centre National de la Recherche Scientifique

Tucker SS, Craine JM, Nippert JB (2011) Physiological drought tolerance and the structuring of tallgrass assemblages. Ecosphere 2(4): 48

Vergutz, L., S. Manzoni, A. Porporato, R.F. Novais, and R.B. Jackson. 2012. A Global Database of Carbon and Nutrient Concentrations of Green and Senesced Leaves. Data set. Available on-line [http://daac.ornl.gov] from Oak Ridge National Laboratory Distributed Active Archive Center, Oak Ridge, Tennessee, U.S.A. http://dx.doi.org/10.3334/ORNLDAAC/1106

Wang, Han; Harrison, Sandy P; Prentice, Iain Colin; Yang, Yanzheng; Bai, Fan; Furstenau Togashi, Henrique; Wang, Meng; Zhou, Shuangxi; Ni, Jian (2017): The China Plant Trait Database. PANGAEA, https://doi.org/10.1594/PANGAEA.871819

Wirth, C. and J. W. Lichstein. 2009. The Imprint of Species Turnover on Old-Growth Forest Carbon Balances - Insights From a Trait-Based Model of Forest Dynamics. Pages 81-113 in C. Wirth, G. Gleixner, and M. Heimann, editors. Old-Growth Forests: Function, Fate and Value. Springer, New York, Berlin, Heidelberg.

Zheng, W. 1983. Silva Sinica: Volume 1-4. China Forestry Publishing House, Beijing.

Österdahl, Sofia(2003): Växtsamhällen på fjällrävslyor iett subarktiskt/alpint landskap / (Plant communities on dens of arctic fox in a subarctic/alpine landscape))
